# Dorsomedial striatal neuroinflammation causes excessive goal-directed action control by disrupting astrocyte function

**DOI:** 10.1101/2024.10.24.620154

**Authors:** Arvie Rodriguez Abiero, Joanne M. Gladding, Jacqueline A. Iredale, Hannah R. Drury, Elizabeth E. Manning, Christopher V. Dayas, Amolika Dhungana, Kiruthika Ganesan, Karly Turner, Serena Becchi, Michael D. Kendig, Christopher Nolan, Bernard Balleine, Alessandro Castorina, Louise Cole, Kelly J. Clemens, Laura A. Bradfield

## Abstract

Compulsive actions are typically thought to reflect the dominance of habits over goal-directed action. To address this, we mimicked the striatal neuroinflammation that is frequently exhibited in individuals with compulsive disorders in rats, by injecting the endotoxin lipopolysaccharide into the posterior dorsomedial striatum, and assessed the consequences for behavioural control. Surprisingly, this manipulation caused rats to acquire and maintain goal-directed actions under conditions that would otherwise produce habits. Immunohistochemical analyses indicated that these behaviours were a result of astrocytic proliferation. To probe this further, we chemogenetically activated the Gi-pathway in striatal astrocytes, which altered the firing properties of nearby medium spiny neurons and modulated goal-directed action control. Together, results show that striatal neuroinflammation is sufficient to bias action selection toward excessive goal-directed control via dysregulated astrocyte function. If translatable, our findings suggest that, contrary to conventional views, individuals with striatal neuroinflammation might be more prone to maladaptive goal-directed actions than habits, and future interventions should aim to restore appropriate action control.

## Introduction

Striatal neuroinflammation is a core neuropathological feature of mental health disorders that feature compulsivity, such as obsessive compulsive disorder (OCD) and substance use disorder (SUD) (*1–4*). Individuals with these disorders perform actions repetitively, often against their desires and despite negative consequences. This has led to the prevailing hypothesis that compulsions arise from a disruption to goal-directed actions and an overreliance on habits (*5–8*). However, a recent article (*9*) challenged this view, suggesting that substance use disorder is more aligned with goal-directed control, an assertion that has prompted much debate within the field (*10–12*). To distinguish between these competing hypotheses, we here attempt reconcile this contradiction at the level of neural mechanism, by investigating how inducing striatal neuroinflammation in rats alters the balance of action control.

The neural circuits of goal-directed and habitual actions have been extensively investigated over the last three decades with considerable homology detected between rodents, primates, and humans (*13–15*). Foundational studies (*10, 11*) revealed that these two types of action control are controlled by distinct, parallel circuits in the cortico-striatal network (although more recent work has called into question how definitive these distinctions might be (*18–20*)). Classically, disrupting one circuit has been shown to shift behaviour to the other action control system, reflecting the behavioural changes seen compulsive disorders. However, the experimental approaches (typically lesions or pharmacological inactivation) do not adequately replicate the subtle neural disturbances observed in these disorders, where widespread neuronal silencing or death is either absent, or present only late in disease progression, long after symptoms appear (*3, 21, 22*). Therefore, the question remains as to what drives this shift in action control. Recent research implicates stress as a common precipitating factor in psychiatric disorders (*20*). At a neural level, stress is almost certainly exerting this effect through neuroinflammation (*23*). Moreover, post-mortem and neuroimaging studies have consistently reported increased markers of neuroinflammation in the striatum in individuals with compulsive disorders (*2, 4, 21, 24–27*). Accordingly, we modelled this physiologically relevant neuropathology in rats to determine the consequences for goal-directed versus habitual action control.

Specifically, we infused the gram-negative bacterial endotoxin and neuroinflammatory mimetic lipopolysaccharide (LPS) into the into the posterior dorsomedial striatum (pDMS) to induce a localised neuroinflammatory response (*28–31*). We then assessed whether rats would show intact action selection across a range of assays probing both cue-guided and free operant choice behaviour. Among these, outcome devaluation procedures provided a particularly crucial test of goal-directed action, as intact performance (i.e. selective responding for a valued outcome over a devalued one) reflects both the sensitivity to outcome value and the contingency between the action and outcome: the two defining features of goal-direction (*13, 32*).

We targeted the pDMS in particular because of its established role as the ‘neuroanatomical locus of goal-directed action’ in rodents (*13, 15*), and its homology to the human caudate nucleus that expresses elevated neuroinflammatory markers in individuals with compulsive disorder (*2–4, 24–27*). Behaviourally, striatal neuroinflammation produced a bias towards excessive goal-directed control and immunohistochemical results suggested a role for astrocytes in this behaviour. Thus, in a final experiment, we chemogeneticially activated hM4Di receptors expressed on pDMS astrocytes during the same behavioural assays to determine whether goal-directed action control depends on intact pDMS astrocytic function.

## Methods and Materials

### Animals and housing conditions

A total of 176 Long-Evans rats, approximately half male, half females, weighing 180–350g and 8-10 weeks of age at the beginning of each experiment were used for this study. Rats were purchased from the Australian Research Centre, Perth, Australia, and housed in groups of 2-3 in transparent amber plastic boxes located in a temperature- and humidity-controlled room with a 12-h light/dark (07:00–19:00 h light) schedule. During behavioural training and testing, animals were food restricted at ~85-95% (8-14g chow per day, see supplement for full details). All procedures were approved by the Ethics Committees of the Garvan Institute of Medical Research, Sydney (AEC 18.34), and Faculty of Science, University of Technology Sydney (ETH21-6657), and the University of Newcastle (A-2020-018).

### Surgery

For neuroinflammation experiments, stereotaxic surgery was performed to infuse LPS (5µg/µL) into the into the pDMS (anteroposterior, −0.2mm; mediolateral, ±2.4mm (male), ±2.3mm (female); and dorsoventral, −4.5mm, relative to bregma) and another cohort of animals received LPS injected into their NAc core (anteroposterior, 1.4mm; mediolateral, ±2.2mm; and dorsoventral, −7.5mm, relative to bregma). For chemogenetic experiments, animals received bilateral injections of 1 µl per hemisphere of AAV-GFAP-hM4Di-mCherry (*Addgene*, item ID 50479-AAV5, titer 7×10^12^ vg/mL) or the control AAV-GFAP104-mCherry (*Addgene*, item ID 58909-AAV5, titer 1×10^13^ vg/mL) at the coordinates for pDMS.

### Behavioural Procedures

Behavioural procedures are described here and shown in Figures 1A and 2A. For full details, please refer to the supplement.

**Figure 1.**
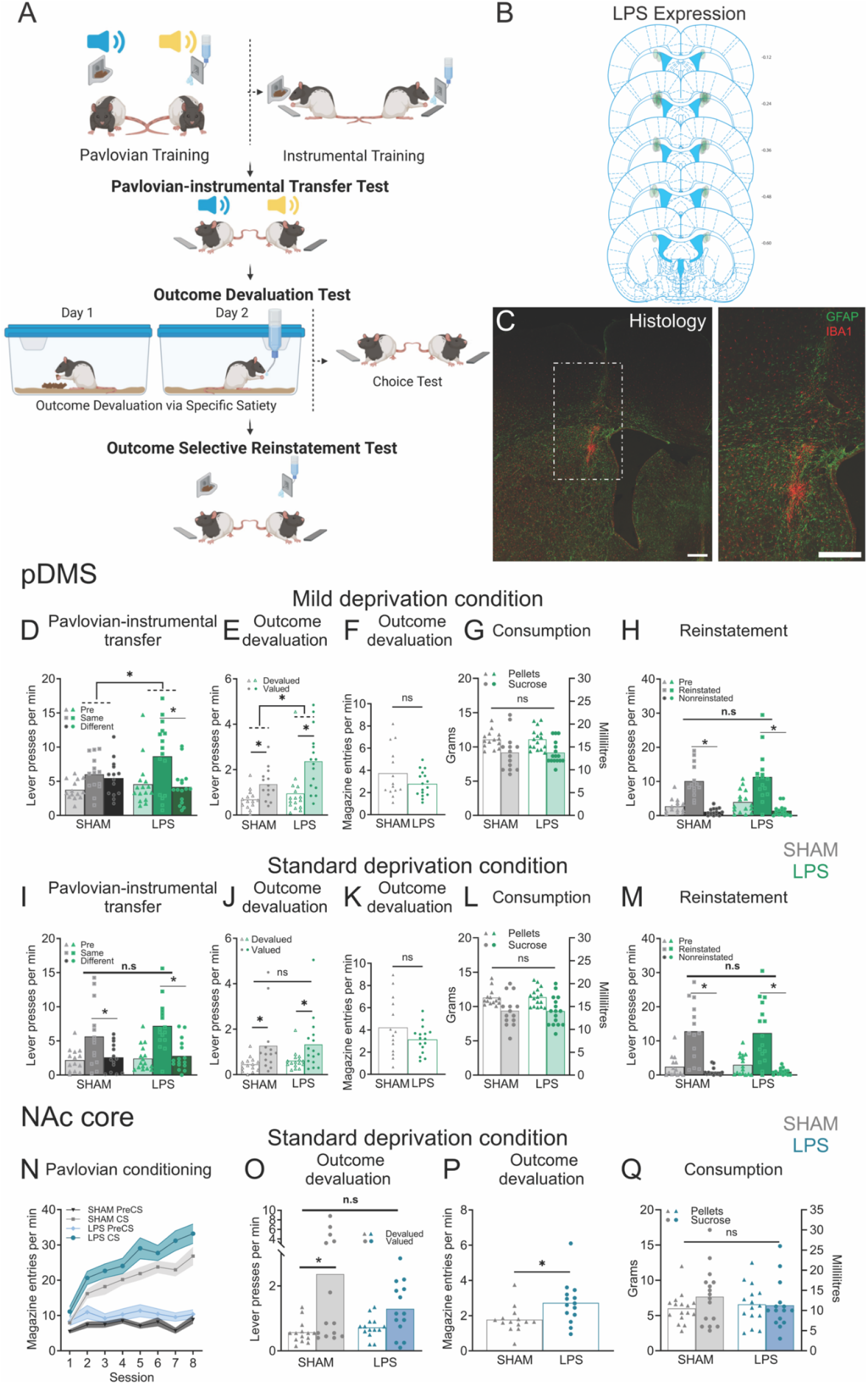
Striatal neuroinflammation causes excessive goal-directed action control in a region specific manner. (A) Experimental procedures for Pavlovian-instrumental transfer, outcome devaluation, and outcome-selective reinstatement, created with *Biorender*. (B) Distribution and locations of the lipopolysaccharide (LPS) injections in the posterior dorsomedial striatum (pDMS) included in the analysis. (C) pDMS image showing LPS placement as labelled with GFAP (glial fibrillary protein) and IBA1 (ionized calcium binding adaptor molecule 1) scale bars = 500 µm, (D-H) Individual data plots and (D,E, & H) mean lever presses (F) magazine entries, or (G) grams/mL consumed during the (D) Pavlovian-instrumental transfer test, (E-F) outcome devaluation test, (G) pre-test feeding and (H) outcome-selective reinstatement test under mild deprivation conditions following pDMS LPS injections, (F) (I-M) Individual data plots and (I, J, & M) mean lever presses, (K) magazine entries, or (L) grams/mL consumed during the (I) Pavlovian-instrumental transfer test, (J-K) outcome devaluation test, (L) consumption, and (M) outcome-selective reinstatement test under standard deprivation conditions following pDMS LPS injections. (N) Magazine entries per min (±SEM) during Pavlovian conditioning, (O) Individual data plots and mean lever presses during the outcome devaluation test, (P) data plots and mean magazine entries during the outcome devaluation test, and (Q) data plots and grams/mL consumed during pre-test feeding following NAc core LPS injections. * Denotes p < 0.05. (pDMS: n = 14 (SHAM), n = 16 (LPS), N = 30; NAc core: n = 14 (SHAM), n = 14 (LPS), N = 28).

### Pavlovian Training

For experiments that involved Pavlovian training, rats were trained once per day for 8 days during which they received eight 2 min presentations of white noise or clicker (4 each) paired with either sucrose solution or pellet delivery.

### Lever Press training

For the initial LPS and the final chemogenetic experiment, rats were trained over 8 days to press left and right levers for sucrose and grain pellets. Lever presses were initially continually reinforced, then progressed to a random ratio schedule. For the experiment employing experimental parameters intended to produce habits, rats received 8 days of two sessions per day during which a single lever was pressed for sucrose, initially on continuous reinforcement, then on random interval schedules.

### Pavlovian Instrumental Transfer test

Each auditory cue (white noise and clicker) was presented four times (8 total) with levers continuously available but no outcomes delivered.

### Outcome Devaluation

For the initial neuroinflammation and the final chemogenetic experiment, outcome devaluation was achieved using specific satiety. For the habit experiment, devaluation was achieved using conditioned taste aversion training. Tests were conducted with either one or both levers present and no outcomes delivered.

### Progressive ratio test

This test was administered during the habit experiment. Animals initially received a sucrose reward for a single lever press, then for 5 lever presses, then n+5 lever presses until breakpoint – with breakpoint defined as 5 min of no lever pressing.

### Electrophysiology

Patch-clamp electrophysiology was performed on DMS tissue 4-6 weeks after local injections with either LPS or adeno-associated viruses (AAVs) resulting in transection with hM4Di-designer receptors exclusively activated by designer drugs (DREADDs). Putative medium spiny neuron (MSN) cell selection was based on MSN cell morphology and post-hoc confirmation of MSN delayed firing action potential (AP) profile, excluding cells without this profile from analysis. Full details are in the supplement.

### Imaging and immunofluorescence analysis

For quantification of glial fibrillary acidic protein (GFAP), ionized calcium binding adaptor molecule 1 (IBA1), and neuron specific nuclear protein (NeuN), a single image was taken of the pDMS and NAc core per hemisphere of each slice (6-10 images in total per brain region of each rat) on a Nikon TiE2 microscope using a 10x objective and Leica STELLARIS 20x air objective for representative images.

### Data and Statistical analysis

Data were collected automatically by Med-PC and uploaded to Microsoft Excel using Med-PC to Excel software. Training data was analysed using two-way repeated measures ANOVAs controlling the per-family error rate at α=0.05. Test data were analysed using complex orthogonal contrasts controlling the per-contrast error rate at α=0.05 according to the procedure described by Hays (*33*). If interactions were detected, follow-up simple effects analyses (α=0.05) were calculated to determine the source of the interaction.

## Results

### Dorsomedial striatal neuroinflammation produced excessive goal-directed action control in rats

Lipopolysaccharide placements in the pDMS are shown in Figure 1B-C. Because we recently showed that neuroinflammation in the hippocampus of mice accelerated this region’s typical function in learning goal-directed actions (*28*) we suspected that neuroinflammation in pDMS might similarly enhance pDMS function to produce ‘excessive’ goal-directed action control, defined as animals exerting such control under conditions for which it is normally absent. Therefore, for the first series of experiments we created conditions to impair action selection in Sham animals by feeding rats a laboratory chow that is relatively high in fat and protein (see supplemental methods and Table 1 for details) on a mild deprivation schedule (approx. 90-95% of their initial body weight) to induce low levels of hunger and arousal (*34*).

**Table 1.** Nutritional information of the lab chow used during the experiment.

|  |  |  |
| --- | --- | --- |
|  | Gordons Specialty Feed | Irradiated Specialty Feed |

| Measurement | LPS | Saline | Significant Difference |
| --- | --- | --- | --- |

|  |  |  |
| --- | --- | --- |
| Protein | 23% | 19% |
| Saturated Fat | 21.3% | 0.78% |
| Mono-unsaturated Fat | 42.9% | 2.06% |
| Poly-unsaturated Fat | 30.7% | 1.88% |
| Crude Fibre | 5% | 5.20% |

**Table 2:**
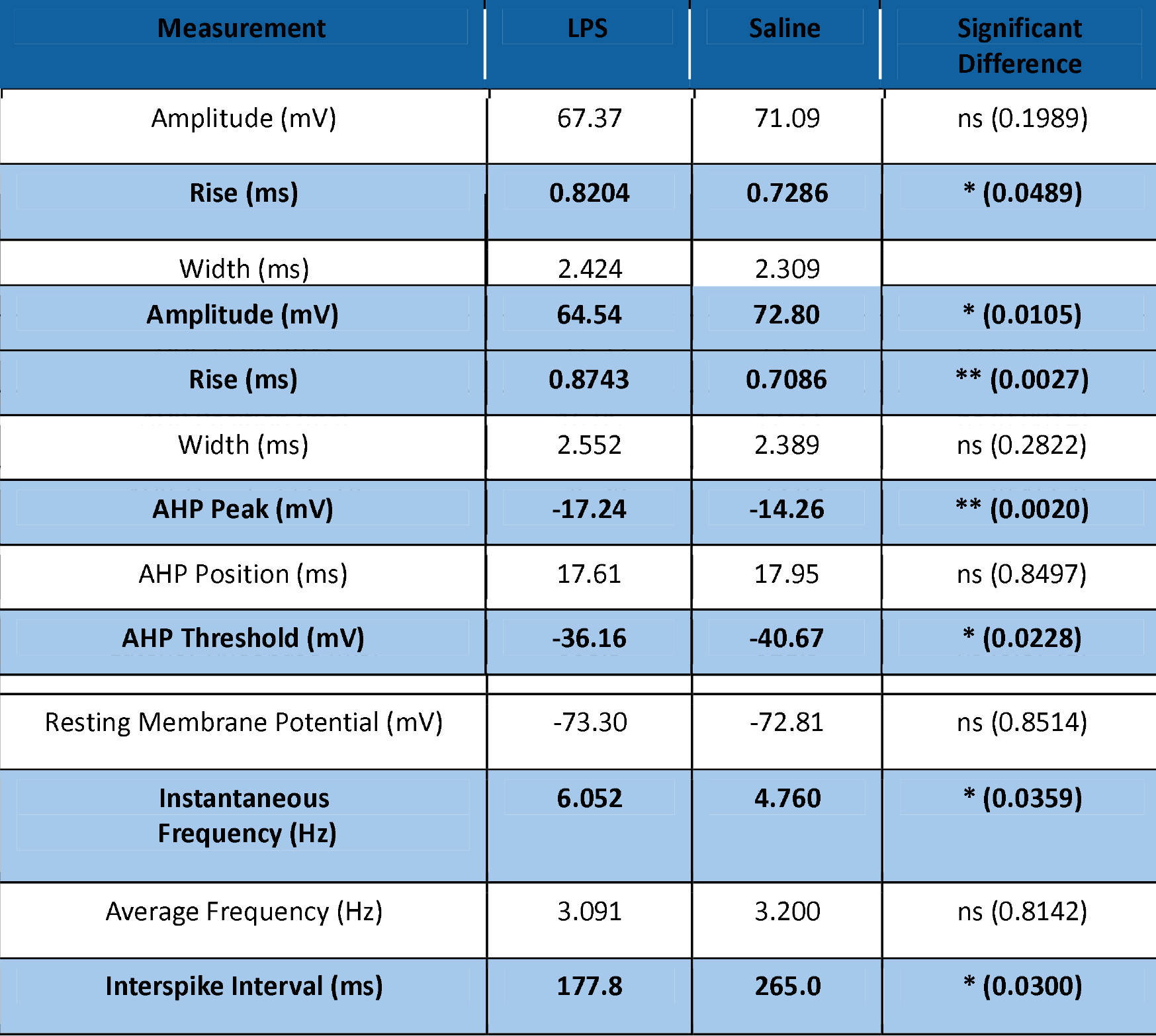
Full results for firing properties of medium spiny neurons in posterior dorsomedial striatum injected with 1ul lipopolysaccharide (LPS, 5mg/mL) at resting membrane potential.

**Table 3:** Full results for firing properties of medium spiny neurons in posterior dorsomedial striatum injected with 1ul lipopolysaccharide (LPS, 5mg/mL) at −80mV.

|  |  |  |  |
| --- | --- | --- | --- |
| Rheobase (pA) | 256.3 | 272.7 | ns (0.6721) |
| Latency Rheobase (ms) | 524.1 | 661.4 | ns (0.0778) |
| Instantaneous Frequency (Hz) | 6.340 | 5.422 | ns (0.3581) |
| Average Frequency (Hz) | 2.545 | 3.067 | ns (0.3126) |
| Interspike Interval (ms) | 238.6 | 230.1 | ns (0.8289) |

**Table 4:** Full results for firing properties of medium spiny neurons adjacent to hM4Di-expressing astrocytes in posterior dorsomedial striatum at resting membrane potential.

| Measurement | ACSF | DREADDs | Significant Difference |
| --- | --- | --- | --- |
| Amplitude (mV) | 75.39 | 75.30 | ns (0.9809) |
| Rise (ms) | 0.7501 | 0.7640 | ns (0.8687) |
| Width (ms) | 2.764 | 3.074 | ns (0.3844) |
| AHP Peak (mV) | -14.95 | -12.07 | ns (0.0879) |
| AHP Position (ms) | 23.59 | 22.64 | ns (0.5459) |
| **AP Threshold (mV)** | **-40.39** | **-45.04** | **\* 0.0118** |
| **Rheobase (pA)** | **185.7** | **125.7** | **\*\* 0.0028** |
| Latency Rheobase (ms) | 640.5 | 441.5 | ns (0.2462) |
| **Resting Membrane Potential (mV)** | **-76.19** | **-64.67** | **\*\* (0.0018)** |
| Instantaneous Frequency (Hz) | 4.762 | 4.749 | ns (0.9910) |
| Average Frequency (Hz) | 3.429 | 2.429 | ns (0.1563) |
| Interspike Interval (ms) | 227.5 | 259.5 | ns (0.5656) |

**Table 5:** Full results for firing properties of medium spiny neurons adjacent to hM4Di-expressing astrocytes in posterior dorsomedial striatum at −80mV.

| Measurement | ACSF | DREADDs | Significant Difference |
| --- | --- | --- | --- |
| Amplitude (mV) | 79.78 | 79.49 | ns (0.9293) |
| Rise (ms) | 0.6602 | 0.6550 | ns (0.8425) |
| Width (ms) | 2.552 | 2.937 | ns (0.1000) |
| AHP Peak (mV) | -13.77 | -11.19 | ns (0.4693) |
| AHP Position (ms) | 21.98 | 19.39 | ns (0.1974) |
| AP Threshold (mV) | -43.85 | -48.04 | ns (0.3269) |
| Rheobase (pA) | 203.3 | 220.0 | ns (0.4859) |
| Latency Rheobase (ms) | 611.2 | 604.2 | ns (0.9769) |
| Instantaneous Frequency (Hz) | 4.478 | 3.664 | ns (0.3537) |
| Average Frequency (Hz) | 3.400 | 2.200 | ns (0.0705) |
| Interspike Interval (ms) | 245.2 | 367.8 | ns (0.4086) |

We first tested cue-guided action selection using specific Pavlovian-instrumental transfer (Fig. 1A). Due to the mild deprivation conditions, we expected transfer to be impaired in controls, as evidenced by equal pressing on each lever regardless of the stimulus presented. However, we expected action selection to be intact in LPS rats despite these conditions, such that they would press more on the lever associated with the outcome predicted by the current stimulus (i.e. the pellet CS would elicit presses on the pellet lever, and likewise for sucrose, Same > Different). This prediction was confirmed (Fig. 1D). Although entries into the food magazine and lever press responses did not differ between Sham and LPS groups during any phase of acquisition (largest F(1,28) = 1.085, p = 0.362, Supp. Fig. 1A-C), on test there was no main effect of group, F < 1, but there was a group x transfer interaction, F(1,28) = 5.710, p = 0.024, driven by a significant simple effect of transfer (Same > Different) for the LPS group, F(1,28) = 15.996, p < 0.001, but not controls (Same = Different), F < 1.

Next, we assessed goal-directed control in the absence of stimuli using outcome devaluation (Fig. 1A). Given the mild deprivation conditions, we predicted that devaluation would be attenuated in Sham controls relative to the LPS group. As predicted, a group x devaluation interaction was observed, F(1,28) = 4.878, p = 0.035, with significant simple effect for both groups that was smaller for group Sham, F(1,28) = 7.445, p = 0.011, and larger for group LPS, F(1,28) = 31.060, p < 0.001 (Fig. 1E). For this test, there was also a main effect of group, F(1,28) = 4.303, p = 0.047, indicating that group LPS responding more overall. Group differences were specific to lever pressing because groups did not differ in prefeeding consumption, F < 1 (Fig. 1G) or magazine entries during test, F < 1 (Fig. 1F).

Following instrumental retraining, rats were tested for outcome-selective reinstatement (Fig. 1A). Because selective reinstatement is not goal-directed (*35*), we expected it to remain unaffected by mild deprivation conditions or pDMS neuroinflammation. This was confirmed, as there was no main effect of group, F < 1, and both groups showed intact reinstatement, i.e. unexpected pellet delivery reinstated responding on the pellet lever, and sucrose delivery likewise reinstated responding on the sucrose lever (Reinstated > Nonreinstated, Fig. 1H), with a main effect of reinstatement, F(1,28) = 67.951, p < 0.001, that did not interact with group, F < 1.

Following this, we explored whether group differences persisted under standard deprivation conditions, (*36–39*) (see supplemental methods for details) for which goal-directed actions should be intact in sham controls and no longer ‘excessive’ in LPS animals. After brief retraining, rats underwent the same transfer, devaluation, and reinstatement tests. This time, performance did not differ between groups on any test: transfer main effect (Same > Different), F(1,28) = 30.605, p < 0.001 (Figure 1I), devaluation main effect (Valued > Devalued), F(1,28) = 12.378, p < 0.001 (Figure 1J), reinstatement main effect (Reinstated > Nonreinstated), F(1,28) = 57.780, p < 0.001 (Figure 1M), with no significant group main effects or interactions, all Fs < 1.

### Neuroinflammation in ventromedial striatum (nucleus accumbens core) preserved instrumental responding, but increased sensitivity to Pavlovian food cues

To test the regional-specificity of neuroinflammation’s effect on goal-directed control, we injected LPS into the NAc core and repeated the same behavioural procedures (*40, 41*). Training and testing were conducted under standard deprivation conditions, based on findings from a pilot study which indicated that NAc core neuroinflammation was unlikely to produce excessive goal-directed control. LPS in the NAc core did not affect instrumental responding during either training or test (Supp. Fig. 1G). Although NAC core neuroinflammation did appear to attenuate devaluation (Fig. 1O) this was not statistically supported as there was no group x lever interaction, F(1,25) = 2.858, p = 0.103). Rather, this attenuation likely resulted from a significant elevation in the competing magazine entry response, group main effect, F(1,26) = 6.02, p = 0.021 (Fig. 1P). LPS rats also made more food magazine entries during Pavlovian conditioning, main effect of group, F(1,25) = 6.962, p = 0.014 (Fig. 1N). Again, these differences were not due to changes in feeding or appetite, because prefeeding consumption did not differ between groups (F < 1, Fig. 1Q).

These results suggest that although NAc core neuroinflammation did not alter instrumental responding, it did enhance Pavlovian responding for food (magazine entries) when response competition from lever pressing was absent (i.e. Pavlovian training) or reduced due to satiety (i.e. devaluation testing). Because enhanced responding to Pavlovian cues has also been claimed to contribute to compulsive-like tendencies (*42, 43*), if translatable, these results suggest that differential distributions of neuroinflammation throughout the striatum could be a multifaceted source of compulsivity.

### Dorsomedial striatal neuroinflammation prevents rats from developing habits

We next wished to confirm that pDMS neuroinflammation could produce excessive goal-directed control under standard deprivation conditions, by preventing habits. We trained a naïve cohort of rats on a single lever using a random interval schedule, as this has been reliably shown to produce habits (*44, 45*); followed by a progressive ratio test to determine whether pDMS neuroinflammation had altered motivation *per se* (Fig. 2A). This was followed by devaluation by lithium chloride injections to induce conditioned taste aversion (Devalued group) whereas the Valued groups received injections of saline. Sham controls were expected to show habitual behaviour (Valued = Devalued) whereas group LPS would remain goal-directed (Valued > Devalued).

**Figure 2.**
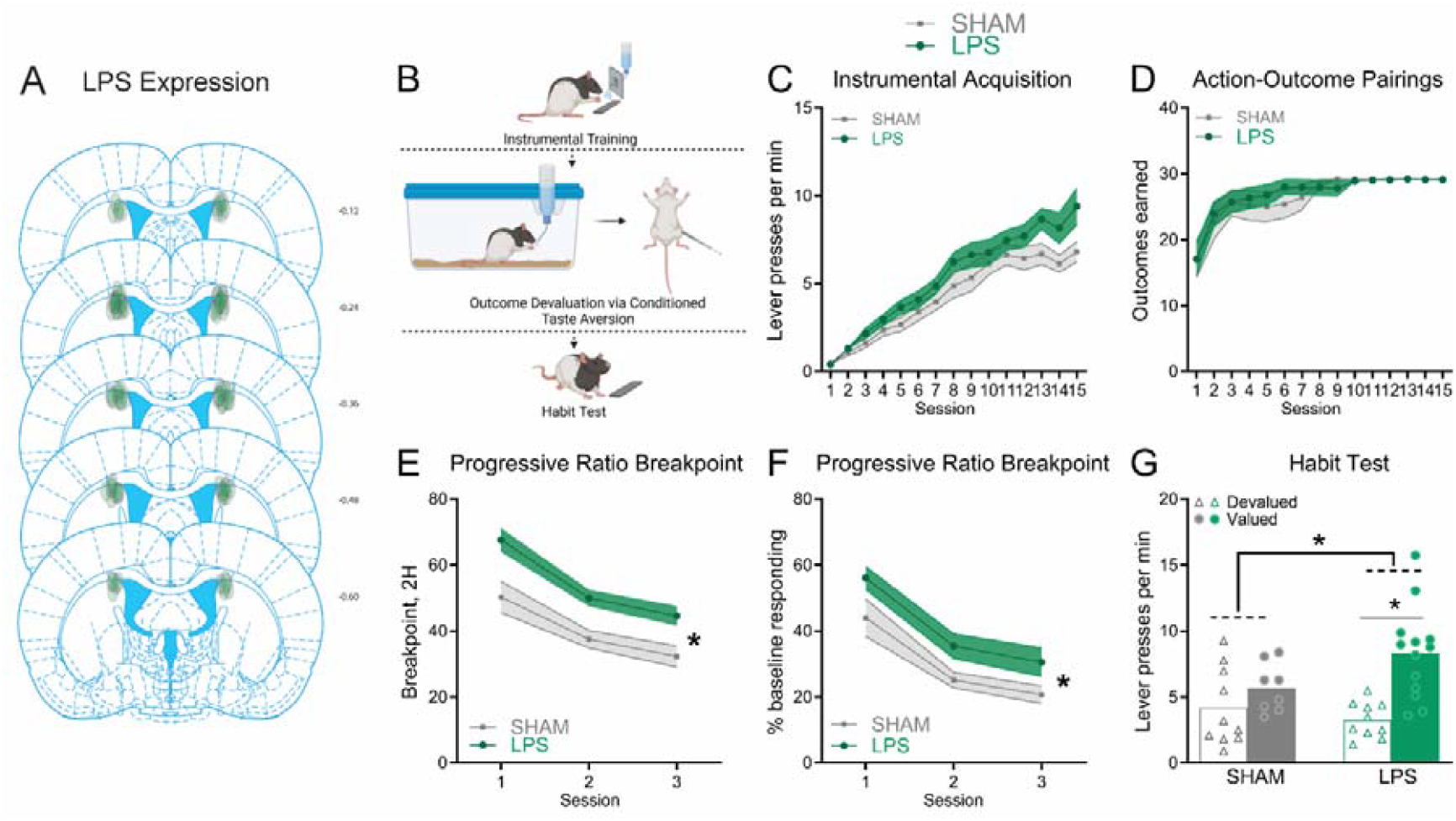
pDMS neuroinflammation prevents the formation of habits. (A) Distribution and localization of lipopolysaccharide (LPS) injections within the posterior dorsomedial striatum (pDMS) included in the analysis. (B) Outcome devaluation procedure designed to promote habits, created with Biorender. (C) Lever pressing per min (±SEM), and (D) Number of action-outcome pairings (±SEM), during instrumental conditioning, (E) Breakpoint (±SEM) obtained during the 2-h, 3-day Progressive Ratio testing schedule, (F) Lever presses during progressive ratio testing (±SEM) presented as a percentage of baseline responding, (G) Individual data plots and mean lever presses during the outcome devaluation habit test. * Denotes p < 0.05. (n = 18 (SHAM), n = 23 (LPS), N = 41).

Groups did not differ on lever press acquisition, though there was a trend toward greater responding in group LPS (Fig. 2B): main effect, F(1,39) = 3.36, p = 0.074. Importantly, the number of action-outcome pairings did not differ between groups, F < 1 (Fig. 2C). LPS rats did have increased breakpoints relative to Shams on progressive ratio testing, however, as there was a main effect of group, F(1,39) = 15.15, p < 0.0014 (Fig. 2D) that remained significant after correcting for baseline press rates, F(1,39) = 6.243, p = 0.0168 (Fig. 2E). As expected, during devaluation testing performance was sensitive to devaluation for LPS rats but not for controls (Fig. 2F). There was no main effect of group, F < 1, but there was a group x devaluation interaction, F(1,37) = 4.373, p = 0.043, comprised of intact devaluation in the LPS group (Valued > Devalued), F(1,37) = 20.198, p < 0.001, and not Shams (Valued = Devalued), F(1,37) = 1.417, p = 0.241. These findings suggest that pDMS neuroinflammation both increased motivation and sustained goal-directed control when controls were habitual.

### Immunohistochemical results indicate a role for astrocytes in excessive goal-directed control

Our final aim was to investigate how pDMS neuroinflammation might cause excessive goal-directed control. To answer this, we first turned to immunohistochemical analyses of tissue from animals who underwent behavioural testing in Figures 1-2 (Figs. 1B and 2B show the regions assessed). For pDMS animals in Figure 1, rats in the LPS group showed significantly higher counts of cells positive for the astrocytic marker GFAP compared to Sham controls, t(28) = 6.26, p < 0.001 (Fig. 3A), and cells positive for the microglial marker IBA1, t(28) = 8.74, p < 0.001 (Fig. 3B), but no significant difference in NeuN-positive cells, a marker of neurons, t(28) = 1.90, p = 0.068 (Fig. 3C). A similar pattern of results was observed in tissue taken from animals in the habit formation experiment (Fig. 2), as shown in Supplementary Figure 3B.

**Figure 3.**
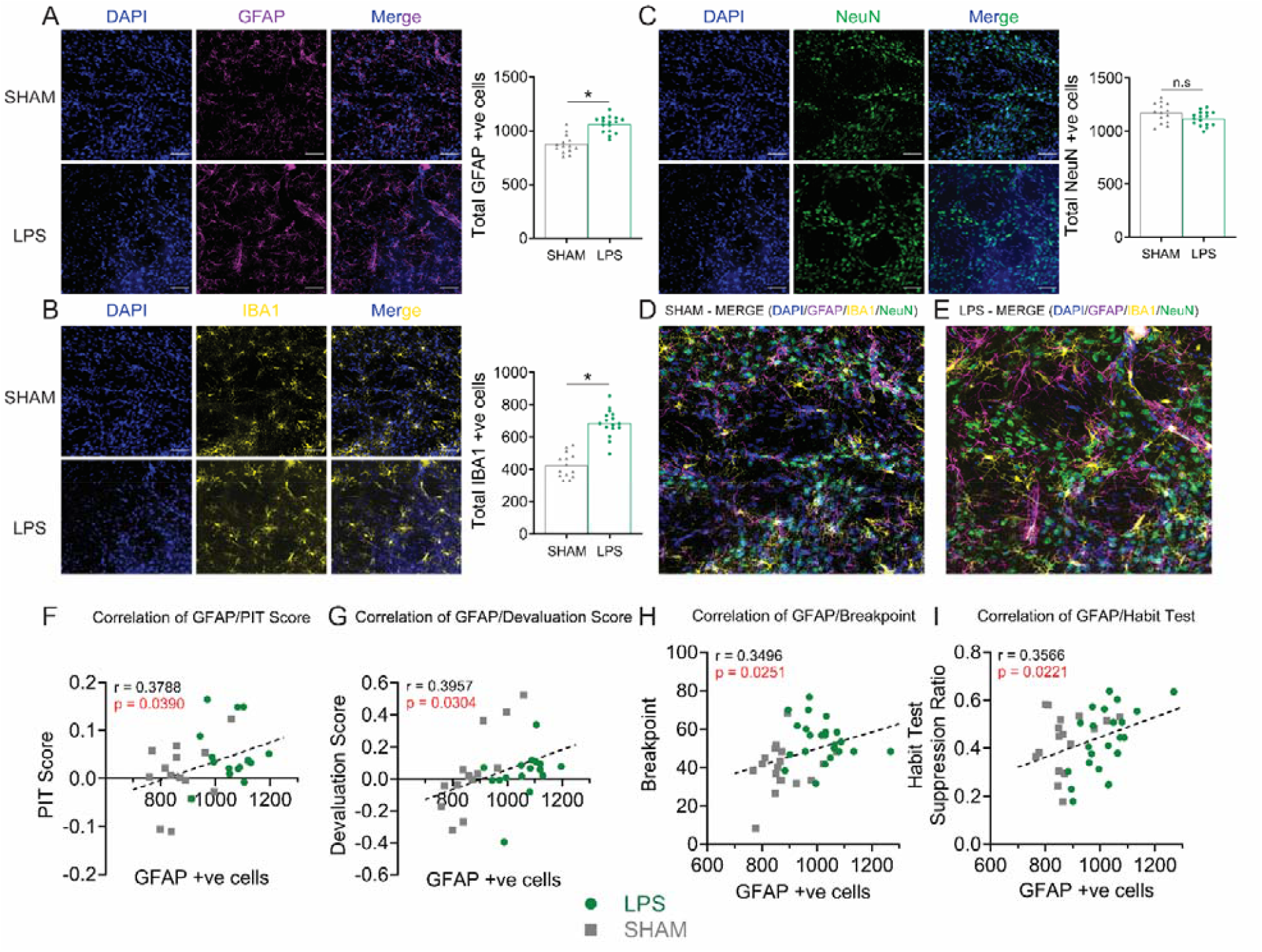
Injections of lipopolysaccharide (LPS) into posterior dorsomedial striatal (pDMS) increased the counts of GFAP and IBA1. Numbers of GFAP+ve cells positively correlated with excessive action control. (A-C) Representative images of pDMS from a Sham (top panel) and LPS-injected rat (bottom panel) immunostained for DAPI and (A) GFAP, (B) IBA1, (C) NeuN, final graphs show individual data points and mean values for quantification of each, (D-E) Representative images of pDMS immunostained DAPI/GFAP/IBA1/NeuN merged from a Sham (D) and LPS-injected (E) rat, (F-I) Correlations between GFAP and behavioural performances. * Denotes p < 0.05, scale bars = 42 µm.

In addition to cell counts, rats in the LPS group also showed elevated signal intensities and other morphological changes in both the astrocyte marker GFAP and microglial marker IBA1 compared to controls (see Supp. Fig. 3 for full results). However, with exception of breakpoint responding that correlated with IBA1 cell counts (Fig. S3C, bottom left), only GFAP measures significantly correlated with action selection on tests where group performances differed (Fig. 3F-I). Importantly, all correlations were calculated using lever press rates normalised to baseline responding, ensuring that observed associations reflect selectivity in behaviour (e.g., Valued > Devalued) rather than general increases in responding. Therefore, while both astrocytic and microglial proliferation were associated with the increase in motivation, only astrocytic proliferation was associated with the enhanced selectivity of actions.

### Neuroinflammation and chemogenetic excitation of Gi-coupled receptors on astrocytes differentially altered the firing properties of adjacent medium spiny neurons

If altered astrocytic functioning underlies the changes in goal-directed actions, it likely does so by altering the activity of nearby neurons, because astrocytes do not have long enough processes to interact with the broader neural circuit of goal-directed control. To explore this, we used *in vitro* whole cell patch clamp electrophysiology to determine how LPS injections in pDMS altered the firing properties of medium spiny neurons (MSNs).

We bilaterally injected LPS or saline into the pDMS of rats, then recorded from acute brain slices 6 weeks later. Recordings were first taken at resting membrane potential (RMP, Fig. S4) then repeated while cells were voltage-clamped at −80mV, consistent with the reported *in vivo* RMP of MSNs (*46*). LPS MSNs displayed a more depolarised AP threshold following a depolarising current steps protocol when voltage-clamped at −80mV (t20.49 = 2.46, p = 0.023, Fig. 4A), whereas no changes were seen for rheobase, instantaneous frequency or interspike interval (Fig. 4B-D). The first AP also showed increased rise time (t37.34 = 3.21, p = 0.003, Fig. 4E, 4H), and decreased amplitude (t31.31 = 2.72, p = 0.011, Fig. 4F, 4H) in LPS MSNs. Furthermore, LPS MSNs showed a significantly more depolarised afterhyperpolarization (AHP) peak (t28.37 = 3.40, p = 0.002, Fig. 4G, 4H). No changes were seen in latency to first spike, half-width, or AHP position (data not shown). These firing patterns suggested that LPS-affected cells in pDMS were less likely to be activated than controls. Taken in conjunction with behavioural results, these findings suggest that LPS in pDMS disrupts the precise excitatory/inhibitory balance necessary for appropriate control over actions, causing goal-directed control to be excessive.

**Figure 4.**
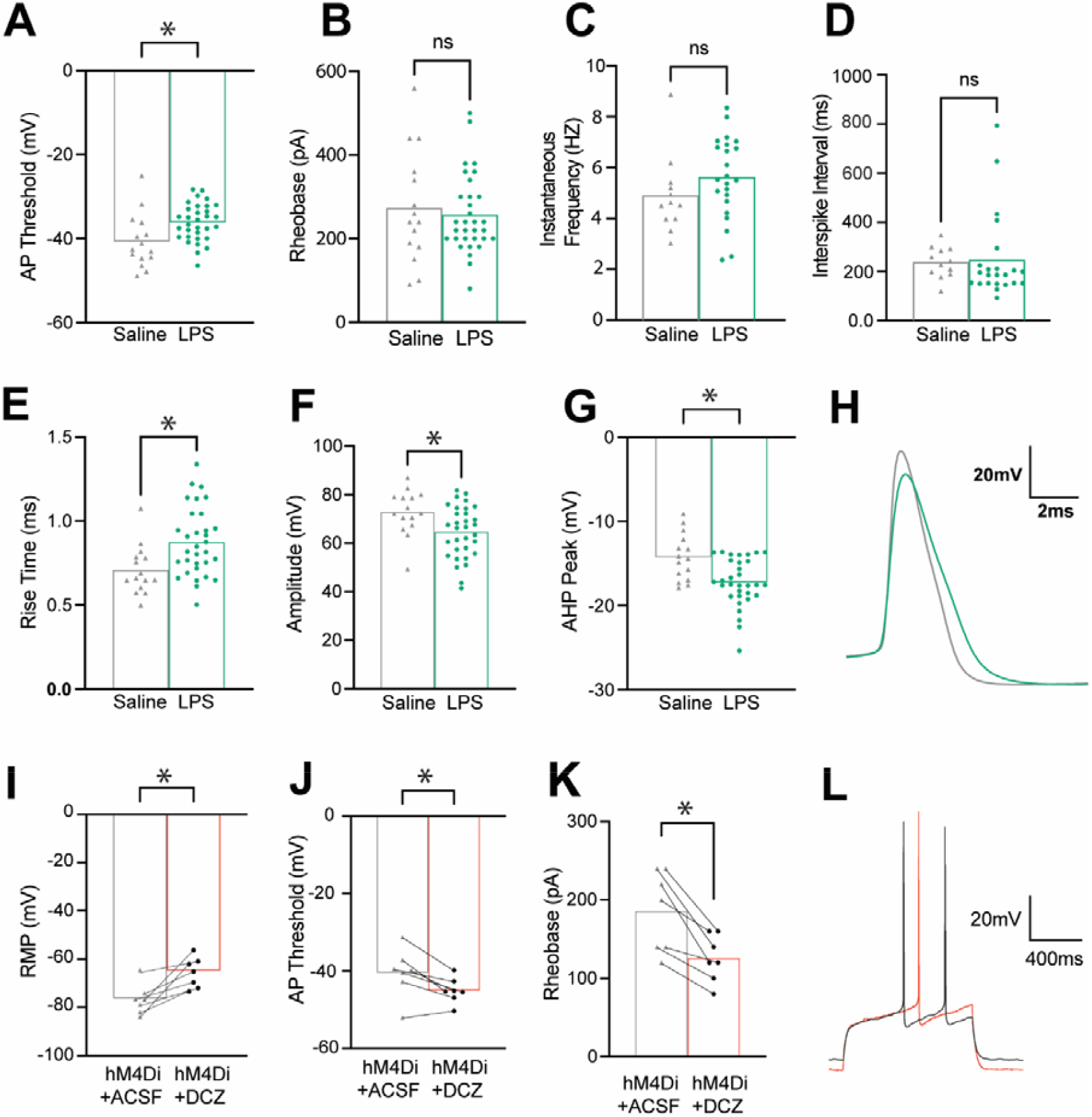
Electrophysiological changes to medium spiny neuron (MSN) action potential (AP) profile and discharge characteristics in pDMS with neuroinflammation or following chemogenetic activation of the Gi-pathway in astrocytes. (A-H) Results of whole-cell patch clamp electrophysiology recordings from MSNs following LPS or sham injections into the pDMS. (A-D) Individual data points showing AP threshold for each MSN voltage clamped at −80mV, (B) rheobase, (C) instantaneous frequency, or (D) interspike interval for each MSN. (E-G) Individual data points for changes to AP profile for each MSN voltage clamped at −80mV including (E) AP rise time, (F) AP amplitude, and (G) an afterhyperpolarisation (AHP) peak. (H) Example cell average trace shows the AP profile characteristics of rise time and amplitude (LPS = green, saline = grey). (I-L) Results of whole-cell patch clamp electrophysiology recordings from MSNs following the application of artificial cerebrospinal fluid (ACSF) then designer receptors exclusively activated by designer drugs (DREADD) agonist deschloroclozapine (DCZ) to astrocytes transfected with hM4Di DREADDs. (I-K) Individual data points showing (I) resting membrane potential (RMP), (J) AP threshold, and (K) rheobase for each MSN. (L) Example cell rheobase traces (ASCF = grey, DCZ = orange). LPS vs saline; LPS at RMP n = 33 cells and at −80 voltage clamp n = 32 cells, from n = 4 animals; saline; n = 15 cells from n = 3 animals. GFAP-HM4Di n = 7 cells from n = 2 animals tested with ACSF then DCZ.

We next manipulated astrocytes specifically. Consistent with the increase in GFAP expression (Fig. 3) a previous study that chemogenetically activated hM3Dq DREADDs on DMS astrocytes observed excessive goal-directed control, albeit in mice undergoing different behavioural procedures (*47*). We therefore aimed to extend these findings by employing a procedure that would reveal how pDMS astrocytes contribute to goal-directed control in their homeostatic form. Based on evidence that Gi-coupled G-protein-coupled receptors (GPCR)s are highly expressed on striatal astrocytes (*48*), and evidence that activating these receptors has been shown to ‘correct’ a number of Huntington-like (*49*) and compulsion-like (*50*) deficits in mice, we used astrocyte-specific hM4Di DREADDs to examine the consequences of Gi pathway activation on neuronal firing and action selection.

Although prior studies have investigated the activation of astrocytic Gi-GPCRs in the striatum (*48, 49*), they have primarily focused on the dorsolateral rather than dorsomedial compartment. This is important, because there are now several studies demonstrating the regional specificity of astrocyte function within the brain, even within the striatum (*48, 49, 51, 52*). Thus, to establish the effects of astrocytic hM4Di activation on neuronal firing properties, we bilaterally injected GFAP-hM4Di-DREADD into the pDMS, and recorded from 7 cells across two animals, firstly in artificial cerebrospinal fluid (ACSF) and then following bath application of hM4Di-DREADD agonist DCZ (1µM). After DCZ application, RMP was significantly more depolarised (t6.00 = 4.14, p = 0.0018, Fig. 4I) shifting cells closer to AP threshold. Then, following the same depolarising current steps protocol used in LPS electrophysiology experiments, AP threshold was lower at RMP (t6.00 = 3.57, p = 0.012, Fig. 4J) further narrowing the range between RMP and AP threshold. Rheobase was also significantly reduced following DCZ application (t6.00 = 4.86, p = 0.003 Fig. 4K,4L). No other changes were seen in AP profile or firing properties (Supp. Fig. 4F-K) with DCZ application at RMP. Recordings were also taken while cells were voltage-clamped at −80mV (Supp. Table 5), blocking the depolarisation of RMP induced by DCZ, and this resulted in no changes to AP profile or firing properties.

This profile of MSN firing contrasts with that produced by LPS injection, and to the results of Kang et al., (*47*) who found that activation of hM3Dq-transfected astrocytes reduced both excitatory and inhibitory postsynaptic potentials (EPSPs and IPSPs) in MSNs. Given that both LPS and hM3Dq activation in astrocytes facilitated goal-directed control whilst producing a distinct profile of neuronal firing, we hypothesised that the activation of Gi receptors on pDMS astrocytes would abolish goal-directed control. Although this may seem counterintuitive in light of prior findings that lesioning or inactivating this structure also abolishes action control (*16, 53*), recent findings paint a more nuanced picture of the conditions necessary for goal-directed action (*54, 55*). In particular, these studies propose that spatially organised neuronal ensembles within the striatum must behave in a precise and complementary manner (referred to as “behavioural syllables”) to produce accurate action selection (*48*). Our electrophysiology results suggest that activating the Gi pathway in pDMS astrocytes disrupts this precision, which we expect to disrupt the behavioural selectivity necessary for goal-directed control.

### Chemogenetic activation of the Gi-pathway in dorsomedial striatal astrocytes abolished goal-directed action control

Above results suggest that the change in astrocyte activity and morphology that occurs as part of the neuroinflammatory response leads to an excessive reliance on goal-directed actions. This implies that the intact signalling of astrocytes in their homeostatic form – i.e. astrocytes that have not undergone a phenotypic shift to a pro-inflammatory-like state – is necessary for intact goal-directed control. To test this idea, behavioural experiments employing chemogenetics were conducted under standard deprivation conditions. Figure 5A and the bottom left panel of Figure 5B show the representative placements of AAV transfection in the pDMS. The bottom panel of Figure 5C shows extensive co-localisation of GFAP and AAV-hM4Di-GFAP-mCherry, and the top panel shows lack of overlap with NeuN, confirming the specificity of transfection for astrocytes.

**Figure 5.**
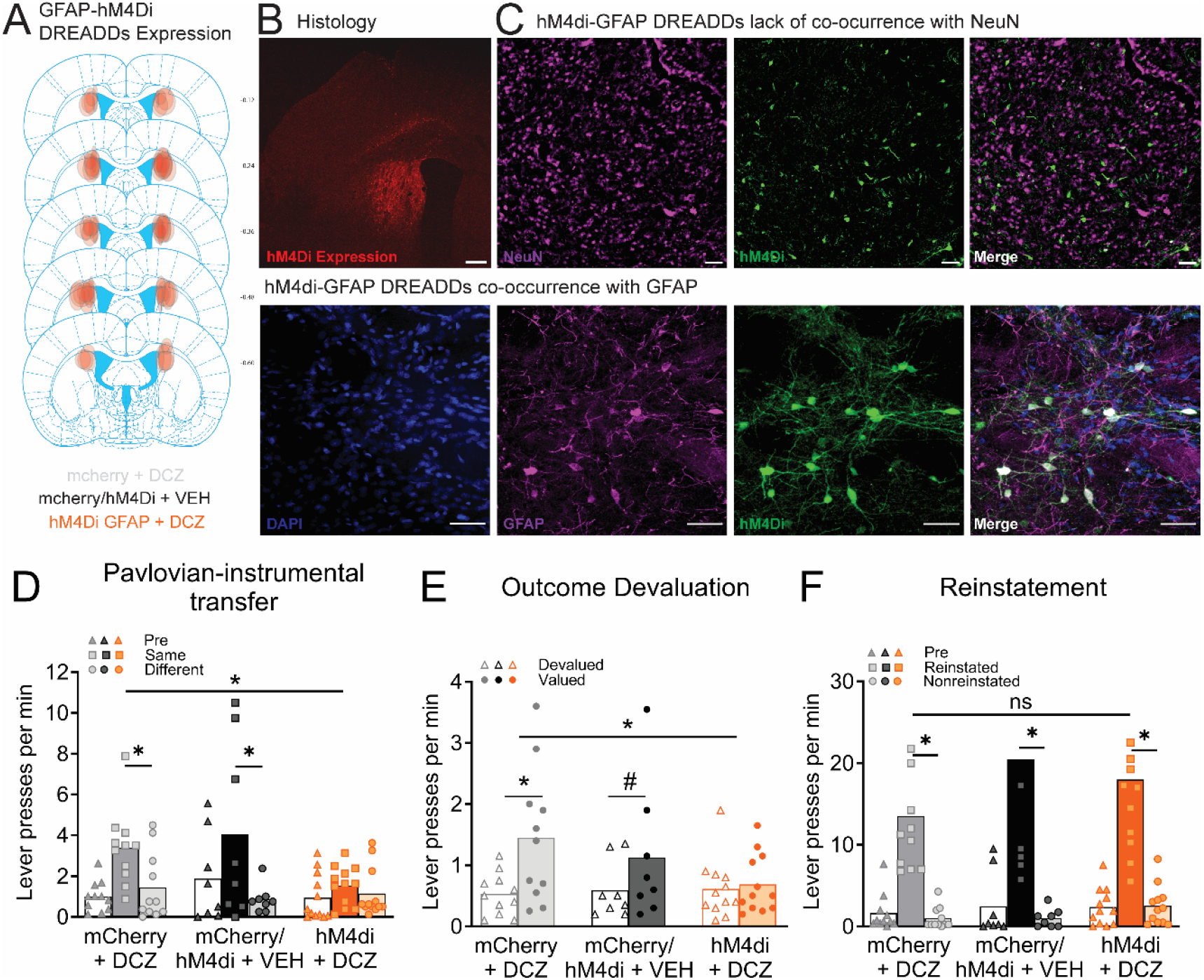
Chemogenetic activation of the Gi-pathway in pDMS astrocytes abolished goal-directed action control. (A) Diagrammatic representation of the distribution and locations of the viral expressions in the pDMS included in the analysis. (B) Histological verification of the GFAP virus expression in pDMS (scale bar = 500 µm). (C) Representative images showing lack of colocalization with NeuN (top panel) and colocalization of mCherry from GFAP-hM4D-Gi-DREADD virus with the GFAP (bottom panel) scale bars = 45 µm. (D-F) Individual data plots and mean lever presses during the (D) Pavlovian-instrumental transfer test, (E) outcome devaluation test, and (F) outcome-selective reinstatement test. * Denotes that the p < 0.05, # denotes p = .055. (n = 8 (hM4Di/mCherry + VEH [hM4Di+Veh =5 + mCherry+Veh n = 3]), n = 11 (mCherry + DCZ), n = 12 (hM4Di + DCZ), N = 31).

Pavlovian and instrumental training were conducted without DCZ administration and proceeded without incident (Supp. Fig. 5A-C, all Fs < 1). However, animals did receive vehicle or DCZ injections to activate the astrocytic Gi pathway 25-30 mins prior to each test. This prevented transfer, which was impaired (Same = Different) in animals that received both the active virus and DCZ (hM4Di+DCZ) but intact (Same > Different) for both vehicle (mCherry or hM4Di+Veh) and DCZ-only (mCherry+DCZ) controls (Fig. 5D). There was a marginal reduction in overall responding, hM4Di+DCZ vs. controls comparison F(1,28) = 3.96, p = 0.056 (comparison between controls, F < 1). More importantly, a group x transfer interaction, F(1,28) = 4.947, p = 0.034, consisting of intact simple effects for groups mCherry+DCZ, F(1,28) = 5.995, p = 0.021, and hM4Di+Veh, F(1,28) = 11.731, p = 0.002, but not group hM4Di + DCZ, F < 1.

It also prevented outcome devaluation, which was intact (Valued > Devalued) for controls but abolished (Valued = Devalued) for group hM4Di+DCZ (Fig. 5D). Specifically, there were no differences in overall responding (both group main effect Fs < 1), but there was a group x devaluation interaction, F(1,28) = 5.494, p = 0.026, driven by intact devaluation simple effects in group hM4Di + Veh, F(1,28) = 16.464, p < 0.001, a marginal simple effect in mCherry + DCZ, F(1,28) = 4.063, p = 0.054, but no effect for group hM4Di + DCZ F < 1.

Finally, performance on selective reinstatement was again intact for all groups; there were no group main effects (largest F was for the comparison between the control groups, F(1,28) = 1.726, p = 0.2), but there was a reinstatement main effect (Reinstated > Nonreinstated) F(1,28) = 67.965, p < 0.001, that did not interact with group F < 1 (Fig. 5E). This demonstrates that the activation of the Gi pathway in pDMS astrocytes does not simply replicate the behavioural results observed following a pDMS lesion or inactivation which have been shown to abolish reinstatement (*16*) as well as devaluation (*16, 53*) and transfer (*56*). Rather, these findings demonstrate a distinct role for astrocytes in regulating the neuronal activity necessary for goal-directed control.

## Discussion

Here we show that striatal neuroinflammation, a common neuropathological feature of compulsive disorders, drives excessive goal-directed action in rats. First, LPS-induced pDMS neuroinflammation promoted such actions under conditions that typically elicit habits. These effects were behaviourally specific, as they did not alter food consumption or selective reinstatement. They were also anatomically-specific: NAc core neuroinflammation did not significantly disrupt instrumental responding. Electrophysiological recordings revealed that, overall, pDMS neuroinflammation reduced the propensity of MSNs to fire, whereas the chemogenetic activation of hM4Di receptors on astrocytes increased MSN firing tendencies. Consistent with these conflicting effects, *in vivo* astrocytic Gi activation disrupted rather than promoted goal-directed actions.

These findings support the emerging hypothesis that individuals with striatal neuroinflammation such as those with compulsive disorders, are acting with cognitive control, albeit inappropriately, under conditions that would otherwise elicit habits (*9, 57*). A potential confound to this interpretation is the observation that rats with striatal neuroinflammation also exhibited higher breakpoints on the progressive ratio test, indicating higher levels of motivation. This could, in principle, explain why the LPS group in the earlier cohort displayed goal-directed control under conditions of low deprivation. However, this account cannot explain why LPS rats also remained goal-directed when Sham rats trained under standard deprivation conditions had transitioned to habitual control (i.e. Fig. 2F). Indeed, increased motivation is typically associated with faster habit formation rather than resistance to it (*58, 59*). Thus, while the while elevated motivation likely contributed to some aspects of performance, it cannot account for the full series of behavioural results. Nevertheless, these findings underscore the complex and sometimes counterintuitive interplay between motivation, habit formation, and goal-directed control.

Although our primary focus was on elucidating neural mechanisms of decision-making, current findings may also offer translational insight into behaviours observed across psychiatric conditions where action control is disrupted. For example, the interpretation of ‘excessive goal-direction’ fits a range of clinical phenomena, including the extreme lengths individuals with substance use disorder undertake to obtain drugs (*60*), or the momentary feeling of relief experienced by individuals with OCD after performing compulsive actions (*61*). It also aligns with observations that individuals with Parkinson’s disease, who also exhibit significant striatal neuroinflammation (*62*), are overly goal-directed, which can slow their responses (*63*). Current findings regarding cue-guided action selection are similarly consistent with enhanced Pavlovian-instrumental transfer effects in rats that have learned to self-administer methamphetamine (*64*), and in humans with alcohol use disorder (*65–67*).

Importantly, we do not claim to model OCD or substance use disorder directly. Rather, we interpret our results as identifying a potential mechanism (striatal neuroinflammation) that may influence decision-making strategies relevant to, but not diagnostic of, these and similar disorders (e.g. Paediatric Autoimmune Neuropsychiatric Disorders Associated with Streptococcal Infections [PANDAS]). If our findings do translate, however, it does bring into question why several lines of evidence suggest that individuals with OCD and substance use disorder over-rely on habits (*5–7*). The following points help reconcile these views. First, individuals with compulsive disorders often show intense focus on a single goal while neglecting competing ones. Thus, if the studies linking compulsivity to habits used goals that weren’t personally salient, participants may have been unmotivated or unable to direct their actions towards them. Second, neuroinflammation is unevenly distributed throughout the brains of individuals with compulsive disorders (*1*) and, as seen here and in past work (*28, 68*), neuroinflammation in different brain regions produces distinct behavioural outcomes. An individual’s dominant behavioural strategy could therefore depend on which brain regions are most affected or could fluctuate dependent on environmental conditions that might preferentially drive cortical, thalamic, and/or nigral/tegmental inputs to different regions. Finally, there is growing evidence that neuronal ensembles within the striatum encode specific sets of action-outcome contingencies (*55, 69, 70*). Depending on the distribution of neuroinflammation and how this interacts with these ensembles, this could promote behaviour that is more goal-directed or more habitual, respectively.

Current results point to astrocytes interacting with these neuronal ensembles to produce goal-directed actions, and it is interesting to consider how this might be achieved. Goal-directed actions are defined by their specificity in achieving a particular goal, such as pressing a lever for a unique outcome (e.g. left lever → pellets, right lever→ sucrose), whereas habits are elicited based on non-specific, prior experience of reinforcement (*71*). A goal-directed response thus requires the selective activation of the neural ensemble that stores the correct response-outcome association, as well as inhibition of the alternate ensemble. Habitual responding does not, instead relying on procedural processes encoded by dorsolateral striatum (*17*). For goal-directed actions, these ensembles consist of the precise, temporally co-ordinated firing of dopamine 1 (D1) and D2-expressing MSNs (*54, 55, 72*). Astrocytes are well-positioned to modulate this activity as they contact both D1 and D2-expressing MSNs to a similar extent (*73*), and express D1 receptors that mediate the dopamine-evoked depression of excitatory neurotransmission (*74*).

Finally, it is worth acknowledging two limitations of our findings. First, although the LPS-induced neuroinflammation in this study had likely progressed past the acute phase by the time of behavioural testing (4-8 weeks post-surgery), it may not fully recapitulate the chronic neuroinflammatory profile experienced by individuals with long-term disorders. This raises the possibility that the behavioural impact of striatal neuroinflammation could shift over time as the neuroinflammatory response evolves (*75*). Second, although current results support a role for homeostatic astrocytic function in goal-directed action control, they do not exclude contributions from other mechanisms, such as the phenotypic responses of microglia. Future studies may wish to address these questions.

In summary, our findings indicate that the alterations to action control experienced by individuals with compulsive disorders are unlikely to be reduced to a single mechanism (*11*), but are multifactorial (*76*), and identify striatal astrocytes as a novel potential therapeutic target to restore adaptive action control. Future research should aim to clarify how different neural and glial mechanisms interact to shape decision-making strategies across contexts and over time.

## Acknowledgements

We thank Thomas Burton for his technical assistance and thank Melissa Sharpe and Octavia Soegyono for helpful discussions regarding the ideas presented in the manuscript. We thank the technical staff at the Garvan Biological Testing Facility at the Garvan Institute of Medical Research and the technical staff at the Ernst Facility at the University of Technology Sydney, and the Bioresearch Facility Staff at the University of Newcastle for technical support. Figures 1A and 2A were created with BioRender.com.

## Author contributions

A.R.A., J.G., S.B., J.A.I., H.R.D., E.E.M., C.V.D., and L.A.B., conceptualised and designed the research, A.R.A., J.G., J.A.I., H.R.D., A.D., K.G., K.M.T., and K.C., performed the research (i.e. data curation), A.R.A., J.G., J.A.I., H.R.D., E.E.M., C.V.D., K.M.T., M.D.K., C.N., L.C., analysed the data, L.A.B., K.M.T., B.W.B., E.E.M., and C.V.D., acquired the funding, A.R.A., J.G., J.A.I., H.R.D., E.E.M., C.V.D., M.D.K., B.W.B., A.C., and L.A.B. wrote the paper (original draft – A.R.A., J.G., and L.A.B., review and editing - A.R.A., J.G., K.M.T, J.A.I., H.R.D., E.E.M., C.V.D., M.D.K., B.W.B., A.C., and L.A.B.).

## Data and materials availability

DOI 10.17605/OSF.IO/297ZU

## License information

**N/A**

## SUPPLEMENTARY MATERIAL

### MATERIALS AND METHODS

#### Experiment 1 and 2: Effects of neuroinflammation in pDMS and NAc core on goal-directed action control

##### Animals and housing conditions

For behavioural experiments, a total of 176 Long-Evans rats [34 for Experiment 1 (15 male and 19 female), 31 for Experiment 2 (15 male, 16 female), 54 for Experiment 3 (27 male and 27 female), 16 for Experiment 4 (6 male, 6 female), and 42 for Experiment 5 (20 male and 22 female)], weighing 180–350 g, 8-10 weeks of age at the beginning of the experiment were purchased from the Australian Research Centre, Perth, Australia, and were housed in groups of 2-3 in transparent amber plastic boxes located in a temperature- and humidity-controlled room with a 12-h light/dark (07:00–19:00 h light) schedule. Experiments were conducted during the light cycle. Before the experiments, all animals were habituated to the laboratory settings for a week with full access to food and water and environmental enrichment which include plastic tunnel, shreds of paper, and wooden object to gnaw. Throughout the training and actual experiment, animals were maintained at ~85% of their free-feeding body weight by restricting their food intake to 8-14g of their maintenance diet per day. All procedures were approved by the Ethics Committees of the Garvan Institute of Medical Research Sydney (AEC 18.34), and Faculty of Science, University of Technology Sydney (ETH21-6657), and the University of Newcastle (A-2020-018).

##### Surgery

Animals were anaesthetized with isoflurane (5% induction, 2–3% maintenance) and positioned in a stereotaxic frame (Kopf Instruments). An incision was made into the scalp to expose the skull surface, and the incisor bar was adjusted to align bregma and lambda on the same horizontal plane. Small holes were drilled into the skull above the appropriate targeted region and animals received bilateral injections by infusing 1 µl per hemisphere of LPS (5ug/ µl) via a 1-µl glass syringe (Hamilton Company) connected to an infusion pump (Pump 11 Elite Nanomite, Harvard Apparatus) into the pDMS (anteroposterior, −0.2mm; mediolateral, ±2.4mm (male), ±2.3mm (female); and dorsoventral, −4.5mm, relative to bregma) and another cohort of animals received LPS injected into their NAc core (anteroposterior, 1.4mm; mediolateral, ±2.2mm; and dorsoventral, −7.5mm, relative to bregma). The infusion was conducted at a rate of 0.15 µl/min, and injectors were left in place for an additional 5 min to ensure adequate diffusion and to minimize LPS spread along the injector tract. The remaining control animals underwent identical procedures but with injection of sterile saline rather than LPS. A nonsteroidal anti-inflammatory/antibiotic agent were administered preoperatively and postoperatively to minimize pain and discomfort. Animals were allowed to recover for 7 days before the onset of any behavioural training.

##### Apparatus

All behavioural procedures took place in twelve identical sound attenuating operant chambers (Med Associates, Inc.,) and these chambers were located within individual cubicles. The ceiling, back wall, and hinged front door of the operant chambers were made of a clear Plexiglas and the side wall were made of grey aluminium. The floor was made of stainless steel grids. Each chamber was equipped with a recessed food magazine, located at the base of one end wall, through which 20% sucrose-10% polycose solution (0.2 ml) and food pellets (45 mg; Bio-Serve, Frenchtown, NJ) could be delivered using a syringe pump and pellet dispenser, into separate compartments respectively. Two retractable levers could be inserted individually on the left and right sides of the magazine. An infrared light situated at the magazine opening was used to detect head entries. Illumination was provided by a 3-W, 24-V houselight situated at the top-centred on the left end wall opposite the magazine provided constant illumination, and an electric fan fixed in the shell enclosure provided background noise (≈70 dB) throughout training and testing. The apparatus was controlled, and the data were recorded using Med-PC IV computer software (Med Associates, Inc.). The boxes also contained a white-noise generator, a sonalert that delivered a 3 kHz tone, and a solanoid that, when activated, delivered a 5 Hz clicker stimulus. All stimuli were adjusted to 80 dB in the presence of background noise of 60 dB provided by a ventilation fan. Outcome devaluation procedures took place in transparent plastic tubs that were smaller, but otherwise identical to the cages in which rats were housed.

##### Food restriction and Chow maintenance

One week following recovery from surgery, animals underwent 3 days of food restriction before the onset of lever press training. During this time animals received 10-14g of chow per day, and their weight was monitored daily to ensure it remained at ~85% of their pre-surgery body weight. For the initial Pavlovian and instrumental training, as well as the first round of testing for sPIT, devaluation, and reinstatement, the chow that rats were maintained on the higher-fat, higher-protein Gordons Specialty Feed (see Table 1). Following reinstatement testing, animals were switched to a lower fat, lower protein Irradiated Specialty Feed’s chow (see Table 1) and re-trained and re-tested.

##### Pavlovian training

For the first 8 days, animals were placed in operant chambers for 60 min during which they received eight 2 min presentations of two conditioned stimuli (CS; white noise or clicker) paired with one of two outcomes (sucrose solution or pellet) presented on a random time schedule around an average of 30 s throughout each CS presentation. Each CS was presented 4 times, with a variable intertrial interval (ITI) that averaged to 5 min. For half the subjects, tone was paired with sucrose and noise with pellets, with the other half receiving the opposite arrangement. Magazine entries throughout the session were recorded and reported for the 2 min prior to each CS presentation (PreCS) and the 2 min during each CS presentation.

##### Lever press training

Following Pavlovian training, animals were trained to press a left and right lever over 8 days which earned the same sucrose and grain pellet outcomes. Specifically, for half of the animals, the left lever earned pellets and the right lever earned sucrose, and the other half received the opposite arrangement (counterbalanced). Each session lasted for 50 minutes and consisted of two 10 minutes sessions on each lever (i.e., four x 10 minutes sessions in total) separated by a 2.5 minutes time-out period in which the levers were retracted and the houselight was switched off. Animals could earn a maximum of 40 sucrose and 40 pellets deliveries within the session. For the first 2 days, animals were trained on a continuous reinforcement schedule (CRF) in which each lever press produced a single outcome. Animals were then shifted to a random ratio-5 schedule for the next 3 days (i.e. each action delivered an outcome with a probability of 0.2), then to a RR-10 schedule (or a probability of 0.1) for the final 3 days. After 40 sucrose solutions and 40 pellets were delivered or 50 minutes had elapsed, whichever came first, the session was terminated, levers were retracted, and house lights switched off.

##### Pavlovian Instrumental Transfer (Specific PIT) test

One day after the end of instrumental training, rats were tested for sPIT performance. For this test, responding on both levers was first extinguished for 8 min to reduce baseline performance. Subsequently, each CS was presented four times over the next 40 min in the following order: clicker-noise-noise-clicker-noise-clicker-clicker-noise. Each CS lasted 2 min and had a fixed ITI of 3 min. Magazine entries and lever pressing rates were recorded throughout the session and responses were separated into PreCS and CS periods (2 min each). Lever presses were recorded but not reinforced.

##### Outcome Devaluation

One day after sPIT testing, rats were given 1 day of instrumental retraining on RR-10 in the manner previously described. On the following day, animals were given free access to either the pellets (20 g placed in a bowl) or the sucrose solution (100 ml in a drinking bottle) for 1 hr. The amount of pellets and sucrose solution consumed each day was measured. Animals were then placed in the operant chamber for a 10 min choice extinction test. During this test, both levers were extended and lever presses recorded, but no outcomes were delivered. The next day, a second devaluation test was administered with the opposite outcome (i.e. if animals were prefed on pellets the previous day they were now prefed on sucrose, and vice versa). Following pre-feeding animals were again placed into the operant chambers for a second 10 min choice extinction test. All test results are reported as averaged across these two tests.

##### Outcome Selective Reinstatement Test

After devaluation testing, rats received one day of instrumental retraining on an RR-10 schedule for 1 day. The next day, animals were tested for outcome-selective reinstatement in which rats received a 15 min period of extinction to reduce baseline performance. They then received four reinstatement trials separated by 4 min each as before, and each reinstatement trial consisted of a single free delivery of either the sucrose solution or the grain pellet presented in the following order: sucrose, pellet, pellet, and sucrose. Responding was measured during the 2 min periods immediately before (pre) and after (post) each delivery.

##### Switching maintenance chow, re-training and re-testing

As noted, a highly palatable home chow (the high-fat/high-protein Gordan’s chow, specifications shown in Table 1) was initially used for Experiment 1 to reduce performance in Sham controls. Following training and testing during which animals were given this chow, rats were switched to a smaller amount (6-8g) of less palatable home chow (lower-fat/lower-protein Specialty Feed’s chow, specifications shown in Table 1) to increase hunger and motivation to lever press for food, with the aim of improving test performance in group Sham.

Following the switch from Gordon’s to Specialty feeds chow, rats were given an additional 4 days of Pavlovian training, and an additional 4 days of intrumental training, then tested for performance on sPIT, outcome devaluation, and outcome-selective reinstatement as before.

##### Tissue preparation

One day after the outcome-selective reinstatement test, animals were sacrificed via CO2 inhalation and perfused transcardially with cold 4% paraformaldehyde in 0.1 M phosphate buffer saline (PBS; pH 7.3-7.5). Brains were rapidly and carefully removed and postfixed in 4% paraformaldehyde overnight and then placed in 30% sucrose. Brains were sectioned coronally at 40 µm through the pDMS and NAc core defined by Paxinos and Watson (2014) using a cryostat (CM3050S, Leica Microsystems) maintained at approximately −20^°^Celsius. The sectioned slices were immediately immersed in cryoprotectant solution and stored in the −20°C freezer.

Later, five representative sections from pDMS and NAc core were selected for each rat. Sections were first washed three times (10 minutes per wash) in PBS to remove any exogenous substances. The sections were then incubated in a blocking solution comprising of 3% Bovine Serum Albumin (BSA) + 0.25% TritonX-100 in 1 x PBS for one hour to permeabilize tissue and block any non-specific binding. Sections were then incubated in anti-GFAP mouse primary antibody (1:300, Cell Signalling Technology Catalog #3670), anti-IBA1 rabbit primary antibody (1:500, FUJIFILM Wako Chemicals U.S.A. Corporation), and anti-NeuN chicken primary antibody (1:1000, GeneTex Catalog #GTX00837) diluted in blocking solution for 72 h at 4°C. Sections were then washed 3 times in 1 × PBS and incubated overnight at 4°C in goat anti-mouse AlexaFluor-488 secondary antibody (1:250, ThermoFisher Catalog #A-11001), donkey anti-rabbit AlexaFluor-568 secondary antibody (1:250, ThermoFisher Catalog #A10042), and goat anti-chicken AlexaFluor-647 secondary antibody (1:250, ThermoFisher Catalog #A-21449), followed by a counterstain with 4⍰,6-diamidino-2-phenylindole (DAPI; Thermo Scientific; 1:1000, diluted in 1x PBS). Finally, every section was mounted onto Superfrost microscope slides (Fisher Scientific) and were coverslipped (Menzel-Glaser) using the mounting agent Vectashield and left to dry overnight in darkness.

##### Imaging and immunofluorescence analysis

For quantification of GFAP, IBA1, and NeuN, a single image was taken of the pDMS and NAc core per hemisphere of each slice (6-10 images in total per brain region of each rat) on a Nikon TiE2 microscope using a 10x objective and Leica STELLARIS 20x air objective for representative images.

##### Microscopy

Images were quantified using imaging software (ImageJ, Fiji Cell Counter), whereby each fluorescent channel was split to isolate and count the cells of interest. Z-stacks were used instead of simply a single image plane. Briefly, the image was adjusted to 8-bit and background subtraction was applied to remove background noise. Thresholding was used to isolate positive stained cells and the threshold for contrast and brightness was adjusted for all images until consistent between images (maximum: 255, minimum: 0). Images were then converted to binary and finally, the Analyze Particles tool was used to quantify the number of cells based on a minimum particle size of 16. ImageJ counted each cell between our parameters and presented it as a “count.” Circularity and perimeter measurement are both part of the Analyze particles plug-in in ImageJ. This was followed by intensity measurement, which is represented as Mean grey value (MGV) and background intensity subtracted from the reported MGV.

##### Data and Statistical analysis

Data were collected automatically by Med-PC and uploaded to Microsoft Excel using Med-PC to Excel software. Pavlovian conditioning and lever press acquisition data was analysed using two-way repeated measures ANOVAs controlling the per-family error rate at α=0.05. If conditions for sphericity were not met, the Greenhouse-Geisser correction was applied. To allow for a more fine-grained analysis of test data, all data for sPIT, outcome devaluation, and outcome-selective reinstatement were analysed using complex orthogonal contrasts controlling the per-contrast error rate at α=0.05 according to the procedure described by Hays (*21*). Acquisition data were expressed as mean ± standard error of the mean (SEM) averaged across counterbalanced conditions. Test data were expressed as individual data points with means. If interactions were detected, follow-up simple effects analyses (α=0.05) were calculated to determine the source of the interaction. For immunohistochemical analysis, counts and intensity were compared between LPS and Sham groups using two tailed t-tests and correlated using GraphPad. Test behaviours were correlated with GFAP, IBA1, and NeuN using the immunohistochemical results from Figure 3. For correlations with behaviour we used a “PIT score”, a “devaluation score”, or a “reinstatement score” that were calculated in such a way as to ensure that any association detected was not driven by baseline differences in lever press responding *per se*, but rather by the animals selectivity of responding for one or the other levers. For these scores, we first calculated suppression ratio (SR) scores on each of the levers individually (i.e., the same and different levers for sPIT, the valued and devalued levers for devaluation, and the reinstated and non-reinstated levers for outcome selective reinstatement) according to Equation 1:

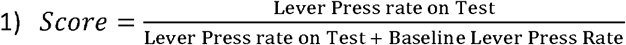

In this equation, “baseline lever press rate” was taken as the average press rate on each lever across the last two days of training prior to test. We then calculated the PIT score by subtracting the normalised scores on the different lever from the normalised scores on the same lever (i.e. Same – Different), such that a higher score indicated better sPIT performance. Likewise, for devaluation we subtracted the normalised scores on the devalued from those on the valued lever (i.e. Valued – Devalued), such that a higher score indicated better devaluation performance, and did the same thing for reinstatement, this time subtracting scores on the nonreinstated from scores on the reinstated lever (i.e. Reinstated – NonReinstated) such that a high score indicated better reinstatement performance. Each of these scores were then separately correlated with GFAP, IBA1, and NeuN counts (correlations with GFAP, IBA1, and NeuN intensity can be found in th excel files uploaded online DOI 10.17605/OSF.IO/297ZU). Values of p < 0.05 were considered statistically significant. The statistical software GraphPad Prism, SPSS, and PSY were used to carry out these analyses.

#### Experiment 3: Effects pDMS neuroinflammation on overtraining-induced habits

##### Surgery

All surgical procedures were conducted identically to that described for Experiment 1.

##### Food restriction and Chow maintenance

For this experiment, animals received only 6-8g of the Irradiated Specialty Feeds chow per day to maintain high motivation conditions. They did not receive Gordon’s chow at any point.

##### Apparatus

All Apparatus were as described for Experiments 1 and 2.

##### Magazine Training

Following recovery from surgery to inject LPS or saline into the pDMS, animals received 3 days of food deprivation and were then given two sessions of magazine training. For these sessions, the house light was turned on at the start of the session and turned off when the session was terminated. No levers were extended. Sucrose solution was delivered at random 60 s intervals for 30 outcomes per session. The session terminated after 45 min or after 30 outcomes had been delivered, whichever came first.

##### Lever Press Training

Following magazine training, animals then received 8 days of instrumental training (two sessions per day) to press a single lever for sucrose solution delivery. Animals received three sessions of continuous reinforcement, four sessions of random interval of 15 s (RI-15), four sessions of RI-30, and four sessions of RI-60. Right and left lever assignment was counterbalanced across animals. Sessions ended, levers retracted and the houselight terminated when 30 reinforcements were earned or after 60 min, which ever came first.

##### Progressive ratio test

Following lever press training, animals underwent 2-h of progressive ratio (PR) testing each day for 3 days. A progressive ratio schedule requires the subject to perform an increasing number of lever presses for the next presentation of a reinforcer (Hodos, 1961). For the current study, the PR was set at n+5. This meant that animals initially received a sucrose reward for a single lever press, then for 5 lever presses, then n+5 lever presses until breakpoint – with breakpoint defined as 5 min of no lever pressing. The number of responses required to obtain each successive delivery of the sucrose reward was collected automatically by Med-PC.

##### Outcome devaluation

The day after progressive ratio testing, animals were given 2 days of instrumental retraining on an RI-60 schedule in the manner previously described. The following day, the sucrose solution was devalued using conditioned taste aversion method for half of the animals. That is, all animals were given ad libitum access to sucrose solution in clear plastic tubs for 30 min each day for 3 days. Immediately after the 30 mins, half of each type of lesion group received an intraperitoneal injection of lithium chloride (0.15 M LiCl, 20 ml/kg) to induce illness which the rat will associate with the outcome, effectively devaluing it, after which they placed back in their home cages. The remaining rats received 0.9% purified saline injections (20 ml/kg) and these animals comprised the valued groups. In total this manipulation yielded 4 groups: Sham-Valued, Sham-Devalued, LPS-Valued, LPS-Devalued. The amount of sucrose solution consumed each day was measured.

##### Extinction test

The day following the last day of LiCl pairings, all animals received a 5 min extinction test. The test began with the insertion of the same lever used during training and ended with the retraction of the lever. Lever presses were recorded, and no sucrose reward was delivered.

##### Tissue Processing and Fluorescent Microscopy

All tissue processing and microscopy were conducted identically to that described for Experiments 1 and 2.

##### Statistical analysis

Lever press and magazine entry data were collected automatically by Med-PC (version 5) and uploaded directly to Microsoft Excel using Med-PC to Excel software. Lever press acquisition and progressive ratio data were analysed using repeated measures (Group x Session) ANOVA controlling the per-family error rate at α=0.05. To allow for a more fine-grained analysis of test data, we used planned, complex orthogonal contrasts controlling the per-contrast error rate at α=0.05 for analyzing the outcome devaluation according to the procedure described by Hays (1973). The amount of sucrose consumed was analysed using Three-Way ANOVA repeated measures. If conditions for sphericity were not met, the Greenhouse-Geisser correction was used. Data analysis was conducted in the manner described for Experiment 3.

#### Experiment 4: Patch-clamp electrophysiology

##### Acute brain slice preparation

To prepare brain slices animals were deeply anesthetised using ketamine injection (100mg/kg, i.p.) and then rapidly decapitated. Following this, brains were rapidly extracted and immersed in ice-cold sucrose substituted artificial cerebrospinal fluid (ACSF) containing (in mM): 236 sucrose, 25 NaHCO_3_, 11 glucose, 2.5 KCL, 1 NaH2PO4, 1 MgCl2, and 2.5 CaCl2. Coronal slices (300µm) of the pDMS were made using a vibrating microtome (VT1200s, Leica, Nussloch, Germany). Slices were then transferred to an incubation chamber containing oxygenated ACSF (120mM NaCL substituted for sucrose) and allowed to equilibrate for 1-hour at room temperature (22-24°C) prior to recording.

##### Patch-clamp electrophysiology

Slices were transferred to a recording chamber and continuously perfused at a rate of 4-6 bath volumes/min with ASCF constantly bubbled with Carbonox (95% O_2_, 5% CO_2_) to achieve a final pH of 7.3-7.4. All recordings were obtained at room temperature (22-24°C), with neurons visualized using near-infrared differential interference contrast optics (IR-DIC). Recordings were restricted to the pDMS in both LPS and hM4Di-DREADD studies, and taken using patch pipettes (4-8 MW, Harvard Glass) filled with a potassium gluconate based internal solution containing (in mM): 135 C_6_H_11_KO_7_, 8 NaCL, 2 Mg_2_-ATP, 10 HEPES, 0.1 EGTA, and 0.3 Na_3_GTP, pH 7.3 (with KOH). Recordings were collected using a Multiclamp 700B amplifier (Molecular Devices, Sunnyvale CA). Signals were sampled at 20kHz, filtered at 10kHz and digitised using an InstraTECH ITC-18 A/D board (HEKA Instruments, Belmore, New York), acquired using Axograph X software (Axograph X, Sydney, Australia). Putative MSN cell selection was based on MSN cell morphology and post-hoc confirmation of MSN delayed firing AP profile, excluding cells without this profile from analysis.

Once whole-cell recording was initiated, series and input resistance were calculated based on the response to a −5 mV voltage step from a holding potential of −70 mV. These measurements were repeated throughout and at the end of all recordings, and data were rejected if this changed by >20 % for an individual cell. Action potential discharge was investigated in current clamp mode, firstly at RMP, and subsequently voltage clamped at −80mV by injection of current when necessary. Depolarising current steps used to evoke AP discharge were increased in 20pA increments for a duration of 1 second and features relating to AP profile were extracted from this data. For recordings using hM4Di-DREADD agonist DCZ (1mM) the above recordings were firstly taken in ACSF and then repeated following bath application of DCZ. Data were analysed offline using Axograph X and Igor Pro 9 (Wavemetrics, Portland, OR) software. AP threshold was taken from rheobase response and AP characteristics were extracted from this including latency to rheobase AP rise time, AP amplitude, AP half-width, AHP peak and AHP position. AP discharge properties were then calculated from rheobase +20pA to determine frequency (mean and instantaneous) and interspike interval.

##### Statistics

Data is presented as mean ± SEM. Unpaired t-tests with Welch’s correction were used to compare LPS and sham affected MSN populations. Paired t-tests were used when comparing unaffected and DCZ treated MSNs.

#### Experiment 5: Chemogenetic activation of Gi-protein-coupled receptors in astrocytes

##### Chemogenetics

The DREADD agonist deschloroclozapine (DCZ) dihydrochloride (NIMH D-925) was acquired from National Institute of Mental Health (NIMH) through the NIMH Chemical Synthesis and Drug Supply Program. DCZ was diluted with normal saline (SAL) (0.9% w/v NaCL) to a final injectable concentration of 0.1 mg/kg (at a volume of 1ml/kg). DCZ was always handled in dim/low light conditions (i.e. a single lamp in a darkened room) and freshly prepared on the morning of each test day.

##### Surgery

All surgical procedures were conducted identically to that described for Experiment 1, except that animals received bilateral injections of 1 µl per hemisphere of AAV-GFAP-hM4Di-mCherry (*Addgene*, item ID 50479-AAV5, titer 7×10^12^ vg/mL). The infusion was conducted at a rate of 0.2 µl/min, and injectors were left in place for an additional 5 min to ensure adequate diffusion and to minimize DREADDs spread along the injector tract. The remaining control animals underwent identical procedures but with injection of AAV-GFAP104-mCherry (*Addgene*, item ID 58909-AAV5, titer 1×10^13^ vg/mL) as control group.

##### Apparatus and Behavioural Procedures

All apparatus and behavioural procedures were conducted identically to that described for Experiment 1, except for outcome devaluation (specific satiety) where animals were given free access to either the pellets or the sucrose solution for 45 mins instead of 1 hr, after which DCZ was administered intraperitoneally (i.p.) and rats returned to their home cage for 25-30 min prior to behavioural testing.

##### Tissue Processing and Fluorescent Microscopy

The extent of the expression was determined using the boundaries defined by Paxinos and Watson (2014). Sections were then stained with Living Colors® DsRed Polyclonal Antibody (1:500, Takara Bio USA, Inc. Catalog #632496) to recognize the mCherry DREADDs expression, anti-GFAP mouse primary antibody (1:300, Cell Signalling Technology Catalog #3670) to check the co-localization, diluted in blocking solution for 72 h at 4°C. Sections were then washed 3 times in 1 × PBS and incubated overnight at 4°C in donkey anti-rabbit AlexaFluor-568 secondary antibody (1:500, ThermoFisher Catalog #A10042), goat anti-mouse AlexaFluor-488 secondary antibody (1:500, ThermoFisher Catalog #A11001), followed by a counterstain with DAPI (Thermo Scientific; 1:1000, diluted in 1x PBS). Sections were mounted and quantified using procedures identical to those described above.

##### Statistical analysis

All statistical analysis was conducted identically as described for Experiment 1.

### Supplemental Figures and Results

**Supplemental Figure 1, relates to Figure 1.**
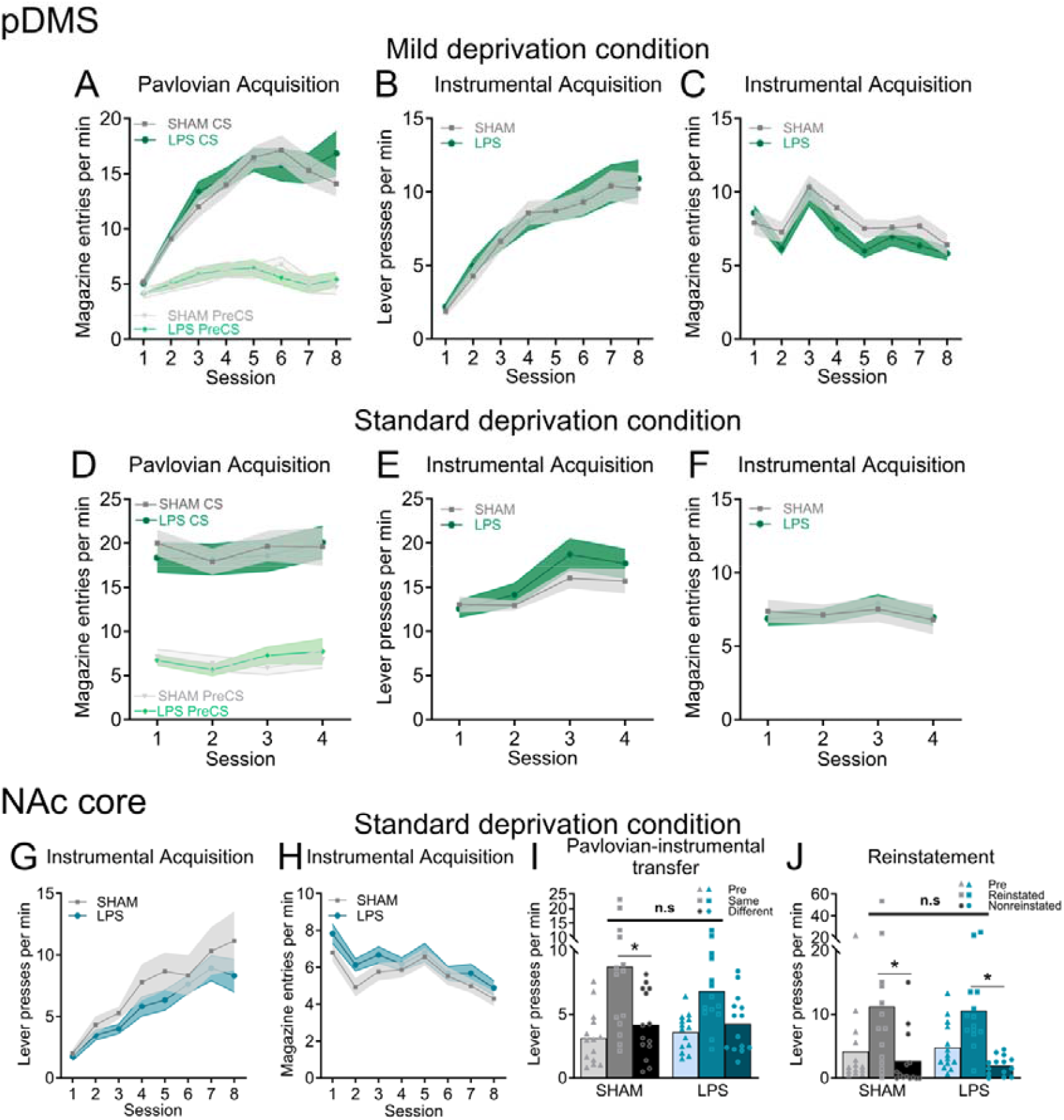
Supplemental behaviour results. There were no significant Sham/LPS differences at any stage of acquisition for pDMS experiment (Experiment 1). (A-C) Acquisition under mild deprivation conditions, (A) Magazine entries per min (±SEM) during Pavlovian conditioning, F(7,196) = 0.669, p = 0.698, for CS x group x session interaction, (B) Lever presses per min (±SEM) during instrumental conditioning, main effect of day F(7,196) = 53.28, p < 0.001, no main effect of group and no day x group interaction, all Fs < 1,(C) Magazine entries per min (±SEM) during instrumental conditioning, main effect of day F(7,196) = 15.282, p < 0.001, no main effect of group and no day x group interaction, all Fs < 1, (D-E) Acquisition under standard deprivation conditions, (D) Magazine entries per min (±SEM) during Pavlovian conditioning, F(3,84) = 2.15, p = 0.111, for CS x group x session interaction, (E) Lever presses per min (±SEM) during instrumental conditioning, main effect of day F(3,84) = 15.87, p < 0.001, no main effect of group and no day x group interaction, all Fs < 1, (F) Magazine entries per min (±SEM) during instrumental conditioning, main effect of day F(3,84) = 2.865, p = 0.041, no main effect of group and no day x group interaction, all Fs < 1. (G-J) Supplemental behavioural results from NAc core neuroinflammation study, there were no significant Sham/LPS differences in instrumental responding for this experiment, (G) Lever presses per min (±SEM) during instrumental conditioning, main effect of day F(7,182) = 27.711, p < 0.001, no main effect of group and no day x group interaction, Fs < 1, (H) Magazine entries per min (±SEM) during instrumental conditioning, main effect of day F(7,182) = 9.692, p < 0.001, no main effect of group and no day x group interaction, Fs < 1, (I) Individual data points and mean magazine entries per min during Pavlovian instrumental transfer testing, a main effect of sPIT, F(1,26) = 14.349, p = .001, that did not interact with the group, F(1,26) = 1.118, p = 0.30. A significant simple effect for the Sham group (Same > Different), F(1,26) = 11.739, p = 0.002, but no such effect (or a marginal simple effect) for the LPS group (Same = Different), F(1,26) = 3.728, p = 0 .064, (J) Individual data points and mean magazine entries per min during outcome selective reinstatement testing, a main effect of reinstatement (Reinstated > Nonreinstated) F(1,26) = 19.278, p < 0.001, which did not interact with any group differences, all Fs < 1.* denotes p < 0.05.

**Supplemental Figure 2, relates to Figure 2.**
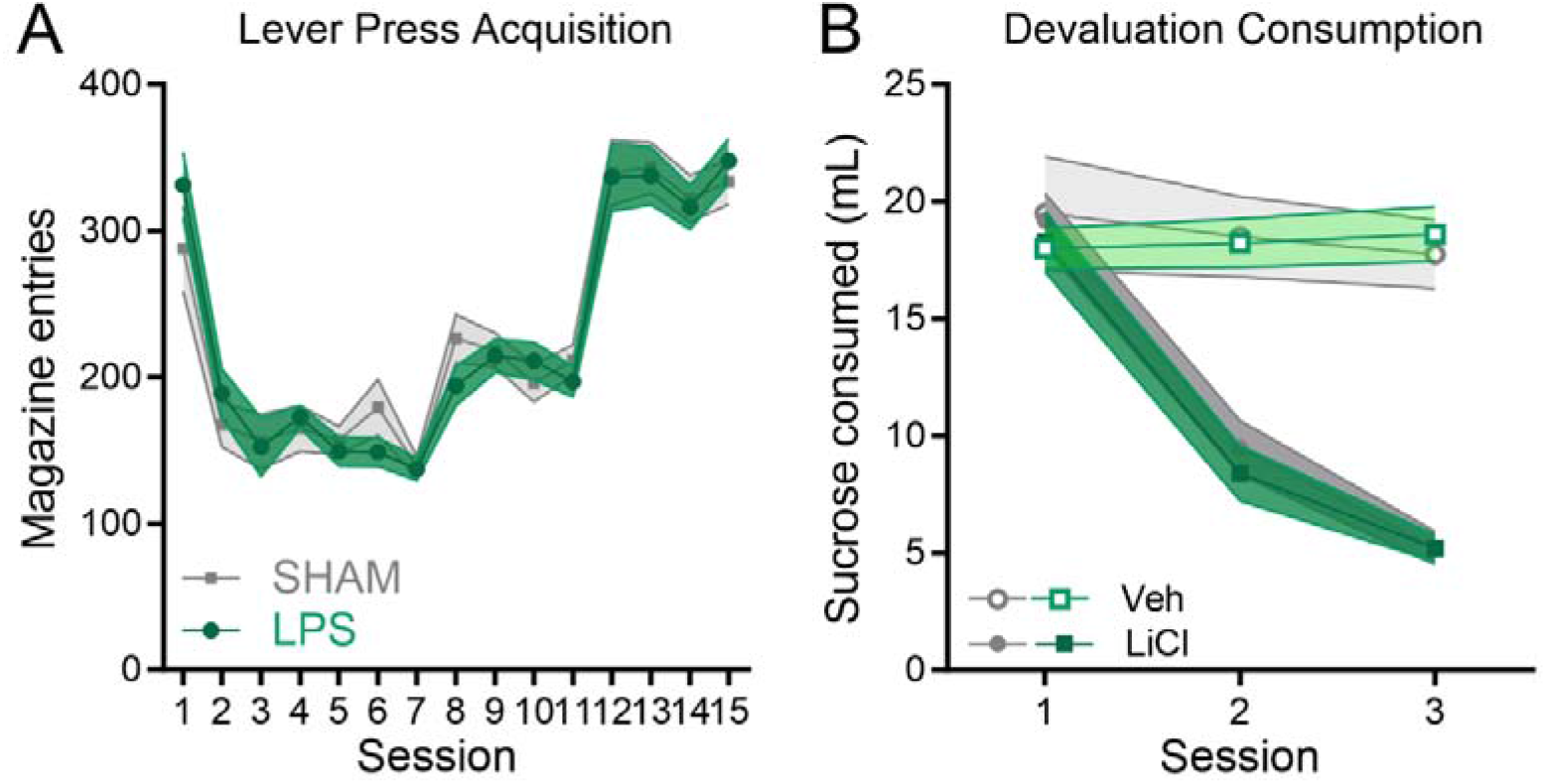
pDMS neuroinflammation did not alter magazine entries during lever press acquisition, nor sucrose devaluation by conditioned taste aversion. (A) Magazine entries per min (±SEM) during instrumental conditioning, main effect of day F(14,546) = 51.27, p < 0.001, no Sham/LPS difference, F(1,39) = 0.002, p = 0.9629, and no day x group interaction, F < 1, (B) Millilitres of sucrose consumed (±SEM) during conditioned taste aversion training, session x devaluation interaction, F(1.64,60.9) = 90.44, p < 0.001 that did not interact with group, F < 1.

**Supplemental Figure 3. Relates to Figure 3.**
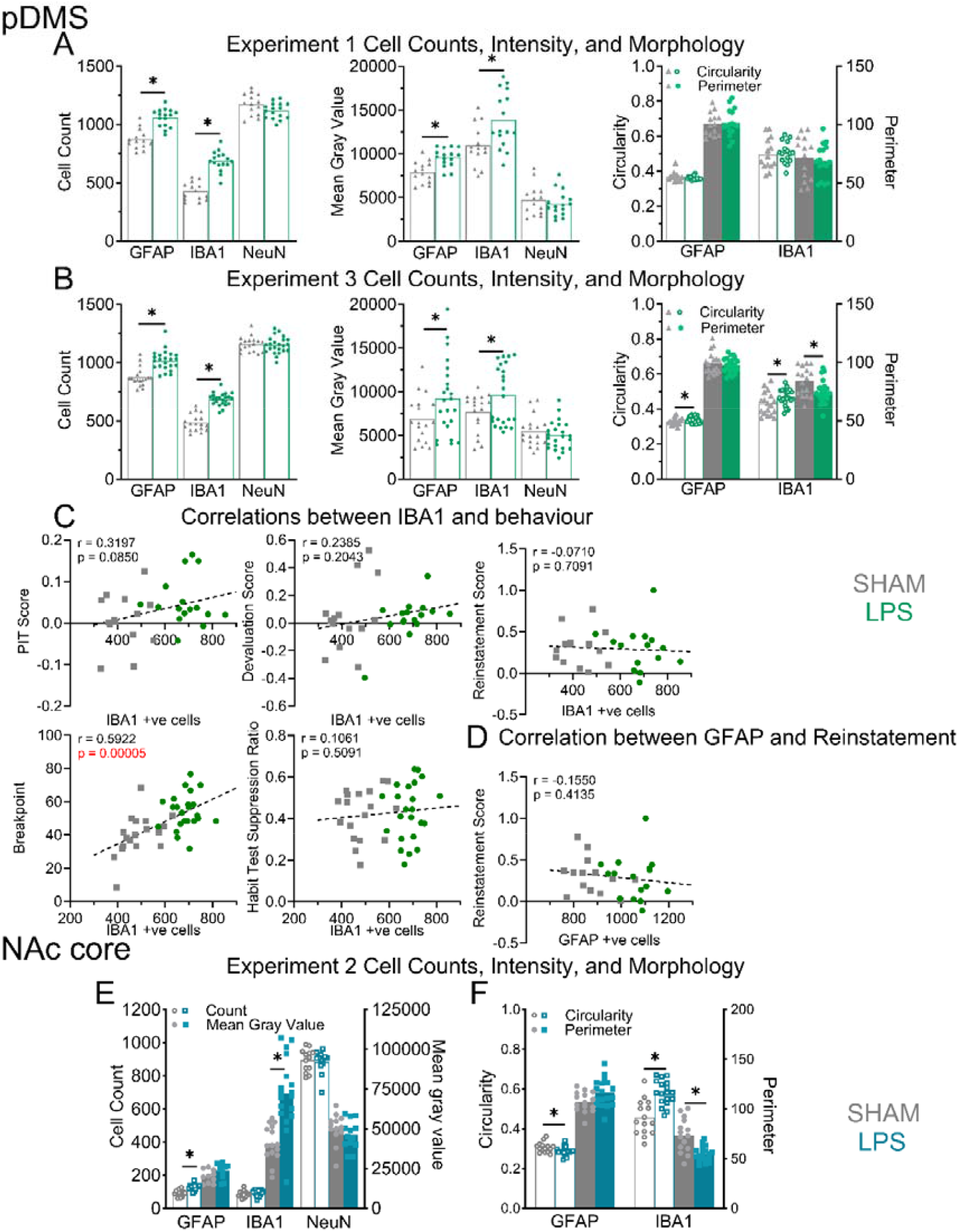
Supplemental immunohistochemical results following injections of lipopolysaccharide (LPS) into posterior dorsomedial striatal (pDMS) in Experiments 1&3. (A-B) Individual data points and mean values for quantification of, from left to right, cell counts, mean gray value, circularity (left y axis), and perimeter (right y axis) of GFAP, IBA1, and NeuN from rats in (A) Experiment 1 and (B) Experiment 3. For Experiment 1: cell counts were significantly higher in LPS tissue relative to Sham for GFAP, t(28) = 6.255, p < 0.001, and IBA1, t(28) = 8.74, p < 0.001, but not for NeuN, t(28) = 1.9, p = .068, mean gray value was likewise significantly higher in LPS tissue relative to Shams for GFAP, t(28) = 4.046, p < 0.0014, IBA1, t(28) = 2.799, p = 0.0092, but not for NeuN, t(28) = 0.7616, p = 0.4527, circularity and perimeter of cells did not differ for any GFAP-positive or IBA1-positive cells in Experiment 1, closest t(28) = 0.77, p = 0.443, for IBA1 perimeter. For Experiment 3: cell counts were significantly higher in LPS tissue relative to Sham for GFAP, t(39) = 5.38, p < 0.001, and IBA1, t(39) = 10.27, p < 0.001, but not for NeuN, t(39) = 0.336, p = 0.074, mean gray value was likewise significantly higher in LPS tissue relative to Shams for GFAP, t(39) = 2.072, p = 0.045, IBA1, t(39) = 2.088, p = 0.043, but not for NeuN, t(39) = 0.8789, p = 0.3848, GFAP-positive cells were significantly more circular for this experiment, t(39) = 2.152, p = 0.0378, as were IBA1-positive cells, t(39) = 2.109, p = 0.0414, whereas perimeter of GFAP-positive cells did not differ between groups, t(39) = 0.7562, p = 0.4541, but was significantly lower in IBA1-positive cells for LPS animals, t(39) = 2.665, p = 0.0113. (C-D) Correlations between (C) IBA1 and (D) GFAP and behavioural performances, r and p values displayed on graphs. (E-F) Results of analyses of immunohistochemical labelling of GFAP, IBA1, and NeuN following NAc core neuroinflammation, (E) Individual data points for cell counts (open shapes and left y axis) and Mean Gray Value (closed shapes and right y axis) values for GFAP (increased counts, t(26) = 4.886, p < 0.001, but not intensity, t(26) = 0.3273, IBA1 (increased intensity, t(26) = 5.110, p < 0.001, but not counts, t(26) = 0.9705, p = 0.5238, and NeuN-positive cells (did not differ between groups), (F) Individual data points for circularity (open shapes and left y axis) and perimeter (closed shapes and right y axis) for GFAP (significant decrease in GFAP circularity, t(26) = 2.412, p = 0.047; perimeter was unchanged, t(26) = 1.955, p = 0.1209) and IBA1 (significant increase in IBA1 circularity, t(26) = 3.829, p = 0.0024; significantly smaller perimeter, t(26) = 3.502, p = 0.0055). * denotes that the p < 0.05.

**Supplement Figure 4. Relates to Figure 4.**
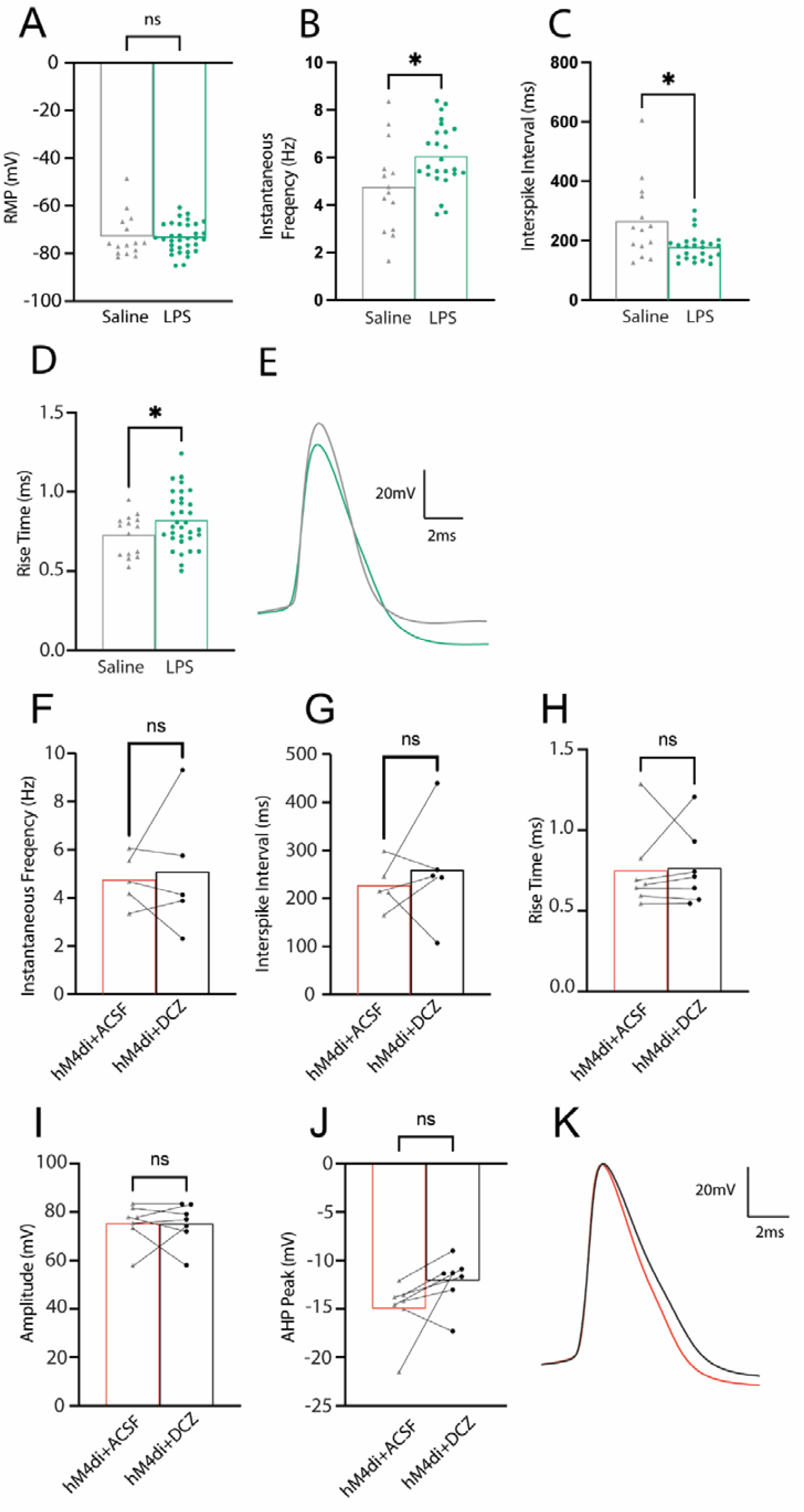
(A-L) Supplemental data from whole-cell patch clamp electrophysiology recordings of MSNs following (A-E) LPS or sham injections into the pDMS or (F-K) following the application of hM4Di-DREADD agonist DCZ to transfected astrocytes. (A-E) Individual data points and means from recordings taken at resting membrane potential (RMP) showing (A) no change to RMP, (B) increased rise time (t36.26 = 2.038, p = 0.0489), (C) increased instantaneous frequency (t20.47 = 2.245, p = 0.0359), and (D) reduced interspike interval (t16.23 = 2.417, p = 0.0278). Rise time changes are further reflected in (E) example cell average traces for the AP profile characteristics (LPS = green, saline = grey). Individual data points and means following DCZ application with MSNs voltage clamped at −80mV (F-K). Data reflected in example cell profile traces presented in (H; ASCF = red, DCZ = black) show AP profile characteristics of rise time and amplitude. LPS vs saline; LPS at RMP n = 33 cells and at −80 voltage clamp n = 32 cells, from n = 4 animals; saline; n = 15 cells from n = 3 animals. GFAP-HM4Di n = 7 cells from n = 2 animals tested with ACSF then DCZ.

**Supplemental Figure 5. Relates to Figure 5.**
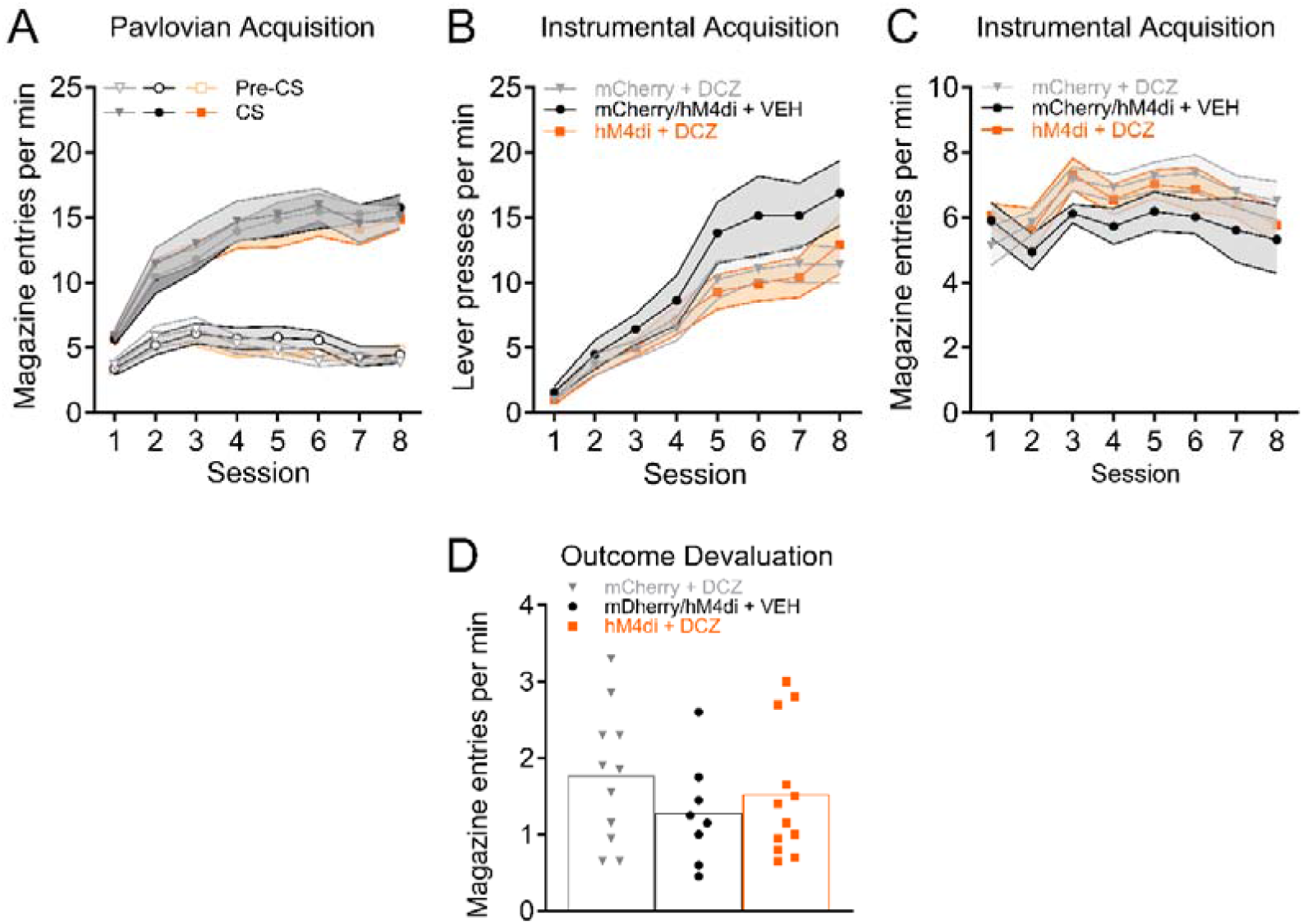
Supplemental behavioural results. (A) Magazine entries per min (±SEM) during Pavlovian conditioning, supported by a main effect of CS period (preCS vs CS) F(1,28) = 742.205, p < 0.001, and of Day F(7,196) = 27.685, p < 0.001, and a Day x CS period interaction (preCS vs CS) F(7,196) = 48.789, p < 0.001. No main effect of group or any interactions with group has been detected, Fs < 1, (B) Lever presses per min (±SEM) during instrumental conditioning, supported by a main effect of day F(7, 196) = 67.262, p < 0.001, no main effect of group (F(2, 28) = 1.803, p = 0.183) and no day x group interaction (F(14, 196) = 1.281, p = 0.222), largest F(5.66, 79.3) = 1.281, p = 0.277, for group x session interaction, (C) Magazine entries per min (±SEM) during instrumental conditioning, supported by a main effect of day F(7, 196) = 8.831, p < 0.001, no main effect of group (F(2, 28) = 1.827, p = 0.180) and no day x group interaction (F(14, 196) = 0.592, p = 0.870), largest F(10.18, 142.47) = 0.592, p = 0.821, for group x session interaction, (D) Individual data points and mean magazine entries per min during outcome devaluation testing, no group differences in magazine entries on test, F(2, 28) = 0.8223, p = 0.4497.

